# Theoretical modeling on CRISPR-coded cell lineages: efficient encoding and optimal reconstruction

**DOI:** 10.1101/538488

**Authors:** Ken Sugino, Jorge Garcia-Marques, Isabel Espinosa-Medina, Tzumin Lee

## Abstract

Delineating cell lineages is a prerequisite for interrogating the genesis of cell types. CRISPR/Cas9 can edit genomic sequence during development which enables to trace cell lineages. Recent studies have demonstrated the feasibility of this idea. However, the optimality of the encoding or reconstruction processes has not been adequately addressed. Here, we surveyed a multitude of reconstruction algorithms and found hierarchical clustering, with a metric based on the number of shared Cas9 edits, delivers the best reconstruction. However, the trackable depth is ultimately limited by the number of available coding units that typically decrease exponentially across cell generations. To overcome this limit, we established two strategies that better sustain the coding capacity. One involves controlling target availability via use of parallel gRNA cascades, whereas the other strategy exploits adjustable Cas9/gRNA editing rates. In summary, we provide a theoretical basis in understanding, designing, and analyzing robust CRISPR barcodes for dense reconstruction of protracted cell lineages.

## Introduction

Genome-wide single-cell molecular profiling has enabled detailed analysis of cell identity and opened an opportunity to trace cell lineage in complex tissues. The identities of individual cells can be inferred from gene expression patterns, via single-cell RNA sequencing. As to cells’ lineage relations, one can cluster related cells based on inheritance of the same somatic mutations acquired by their common precursors. However, unlike single-cell transcriptomes which can be readily verified, the retrospectively built cell phylogeny is only testable in a deterministic lineage that has already been mapped. Since scientists aim to map previously uncharacterized lineages, it is critical to know both the optimal means for reconstruction and the limitations of the methodology. Theoretical modeling is therefore an essential step in designing an ideal lineage tracing experiment.

Cell lineage reconstruction is guided by the developmentally acquired mutations that are retrieved from single cell sequencing. Therefore, the number and diversity of mutations limits the scope and resolution of the cell lineages that can be built. Although the genome can carry innumerable spontaneous mutations [1], sequencing each nucleotide of every sampled cell is not only technically challenging but also financially unrealistic. By contrast, the CRISPR technology is based on gRNA-directed mutagenesis by Cas9 and allows for generation of “genetic barcodes”. These barcodes consist of well-defined units containing gRNA target sequences that one can assess with specific primers. Versatile designs are possible with such CRISPR-based genetic barcodes. First, the virtually infinite gRNA variants permit recruitment of diverse endogenous and/or exogenous sequences into the barcode. Second, a single gRNA target spans only 23 base pairs, thus a densely packed coding block can carry 40 targets per kb. Third, the CRISPR-induced mutations within a given gRNA target site or across adjacent sites are largely predictable and thus programmable. Fourth, there exist variant Cas9-related enzymes that can elicit distinct mutations via cutting the DNA or modifying specific nucleotides. Finally, one can restrict Cas9 activity to cycling cells to prevent post-mitotic mutagenesis, which would be extremely helpful/important for lineage reconstruction. Moreover, for targeted cell lineage analysis, we can also drive Cas9 expression only in the lineage(s) of interest. The enormous versatility of Cas9 plus the superior programmability and trackability of genetic barcodes make CRISPR a uniquely powerful tool for deciphering cell lineages. Establishing a sophisticated system for modeling CRISPR-coded cell lineages would expedite the exploration of creative CRISPR barcode designs. Modeling is the ideal first step to mapping complex cell lineages with near single-cell-division resolution.

As in the phylogenetic analysis of biological species, the task for reconstructing cell lineage is to build ‘trees’ where leaves represent the examined objects (i.e. sampled cells) and the branching patterns outline the implicated developmental history. Because of these similarities, phylogenetic methods are often employed for reconstructing cell lineage trees. The key elements in reconstruction are internal nodes that represent the precursors that give rise to differentially marked descendants. The strategies for estimating those internal nodes therefore govern how to derive a tree that best describes the phylogenetic relationships of leaves. The underlying assumptions in deriving these strategies, however, are different between evolutionary phylogeny and CRISPR-coded cell lineage tracking. For example, the number of possible sites of phylogenetic mutations is often regarded as infinite, whereas the number of CRISPR-coding target sites is limited to those in the barcodes. CRISPR-based mutations are often fixed once mutated (with the exception of homing CRISPR [2]), whereas evolutionary mutations can be overwritten by succeeding mutations. Given these differences, the CRISPR-based method requires independent investigation of optimal reconstruction methods and of the intrinsic limitations of the encoding process.

Knowing the general topology of the underlying cell lineages is also critical for informative cell phylogeny studies. There are two basic modes of cell division. These result in daughter cells with either distinct (asymmetric) or equivalent (symmetric) potentials. Via asymmetric cell division, a stem cell can produce an intermediate precursor which can either divide once into two often distinct post-mitotic cells (type 1) or can itself divide asymmetrically yielding a secondary series of precursors with further restricted potential (type 2). Some long-lived stem cells (e.g. Drosophila neuroblasts) can undergo many rounds of self-renewing asymmetric cell divisions and thus produce an extended cell lineage. By contrast, a developing tissue may expand exponentially through symmetric cell divisions of existing cells (type 3). Distinct challenges exist in tracking lineages across different topologies. Protracted stem-cell-type lineages across many generations require sustained encoding. Whereas, cells that undergo symmetric expansion can carry more coincident mutations. A versatile genetic system for cell phylogeny studies should work robustly for mapping diverse types of cell lineages. Pre-validation in silico is essential to ensure the versatility of any system.

Here, we used theoretical modeling and computer simulation to explore the generation of CRISPR barcodes as well as the algorithms for reconstructing CRISPR-coded cell lineages. We began with modeling the dynamics of the CRISPR-editing assuming a constant Cas9 editing rate across targets and cells. This assumption leads to an exponential reduction in the number of new Cas9 edits along the lineage depth (generation), which therefore severely constrains the trackable lineage depth. We calculated the number of encoding units (Cas9 target sites) minimally required for faithfully tracking every cell cycle in an extended cell lineage. The number of necessary encoding units depends upon the Cas9 editing rate. We found that the optimal editing rate for tracking a series of N cell cycles is simply 1/N. However, even with an optimal editing rate, the number of units needed for tracking protracted Drosophila neuronal lineages (~100 cell cycles per lineage) would be extremely daunting. To enhance the depth of coverage, we explored additional strategies for sustaining the coding capacity beyond the limit set by the exponential reduction. We conceived of two plausible systems that deliver just enough new edits per cell cycle for an entire lineage and thus greatly improve the barcoding efficiency. The first method depends upon making different gRNAs available over time to eliminate early depletion of encoding units. To accomplish this, we propose gRNA cascades. The other solution depends on adjustable Cas9 editing rates, as the desired editing rate increases as the number of available targets decreases.

In search for an optimal tree-building method, we evaluated the depth-dependent error rates in a variety of reconstruction algorithms. We examined those previously employed for CRISPR-based lineage tracking as well as a multitude of different hierarchical clustering methods. Intriguingly, the most faithful tree building was achieved through hierarchical clustering with a distance metric based on the shared edits (an expansion of Russell-Rao metric to non-binary cases) and the complete linkage method. We found that the distance matrix derived from shared edits contains the lineage depth information in a way best suited for reconstructing the lineage in the proper order. In conclusion, The knowledge gained through our theoretical modeling and computer simulation will facilitate dense reconstruction of cell lineages, essential for comprehensive analysis of organism development.

## Results

### Dynamics of CRISPR barcode editing across cell generations

To understand the dynamics of the CRISPR barcode editing, we model the process of editing *nU* units of targets where the rate of editing per cell division is *r*. We assume the editing rate is equal and independent across units and cell division (Fig.1A). Unlike traditional phylogeny, once a unit is edited, it cannot be edited again. Hence homing CRISPR encoding is excluded from the current model. We also denote the number of editing outcome as *nL* (number of levels/alleles), which have equal probability (*p* = 1/*nL*) of being chosen at each editing event. Most of these assumptions are simplified. For example, allele choice is likely to be biased and editing rate could vary between targets. However, we began with this simplified model to delineate the basic dynamics of CRISPR-barcoding with relatively simple mathematics.

**Figure 1:**
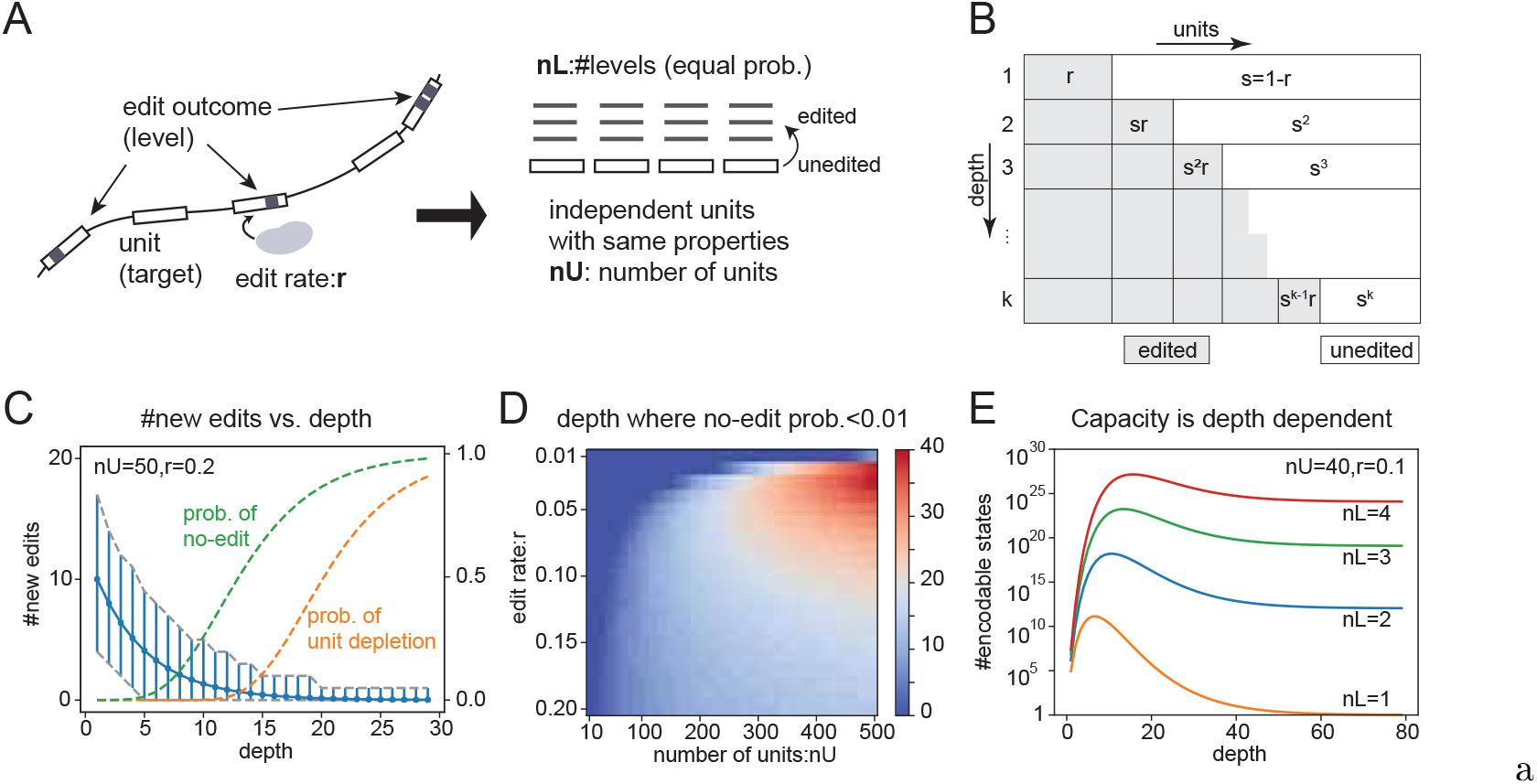
Dynamics of encoding. (A) Schematics of the model of the CRISPR encoding process. Critical prameters: *nU*: number of encoding units, *nL*: number of edit outcomes, *r*: edit rate/division. (B) Schematics showing how average numbers of edited units transition as cells divide. *s* = 1 − *r*. (C) Average (*nUrs*^*k*−1^) and 99% range of new edits (calculated from binomial distribution *B*(*nU,rs*^*k*−1^) at each cell cycle (blue lines), probability of no-edit (green, (1 − *rs*^*k*−1^)^*nU*^ and unit depletion (orange, (1 − *s^k^*)^*nU*^), *nU* = 50 and *r* = 0.2. (D) Heatmap showing the maximum depth where probability of no-edit is smaller than 0.01 for ranges of edit rates (y-axis: 0.01 to 0.20) and number of units (x-axis: 10 to 500). (*z* = *argmax_k_*((1 − *rs*^*k*−1^)^*nU*^ < 0.01), maximum depth where *P*(*noedit*) < 0.01, color-coded). (E) Capacity vs. depth for *nL* =1 ~ 4. Capacity is the number of encodable states, and equals _*nU*_*C_ne_nL^ne^* where ne is the average number of edited units and at depth *k*:*ne*(*k*) = *nU*(1 − *s^k^*).

Having a certain number of new edits at each cell cycle is critical for high-resolution reconstruction of a lineage. We call the number of cell divisions experienced by a cell depth. At given depth, under the above assumptions, the number of newly edited units follows a binomial distribution with the success probability as *r* and the number of trials being the number of remaining unedited units. On average, the number of remaining unedited units decreases exponentially: *nU*(1 − *r*)^*k*−1^ for a depth of *k*; thus the average expected number of newly edited units also drops exponentially: *nUr*(1 − *r*)^*k*−1^ (Fig.1B, C). This phenomenon suggests that the ability to tell ancestral relationship in a lineage also rapidly decreases.

To record contiguous cell cycles along the lineage depth, we need to know the distribution of the number of new edits at each depth. Since the convolution of binomial distribution is again binomial, this distribution can be easily calculated (see supplement) and the 99% range of the number of new edits at each depth is shown in Fig.1C (vertical blue lines). One noticeable finding from this plot (other than exponential decay) is that the occurrence of “no-edit” (i.e. #now-edits 0) happens well before all the units are edited (Fig.1C green and orange lines). This indicates that encoding failure (i.e. cannot distinguish parent-children) happens much earlier than depletion of unedited units. Fig.1D shows the maximum depth where probability of no-edit is less than 0.01 for a range of edit rates and of total units. For a fixed edit rate, one needs to increase the number of units to sustain the encoding over a larger depth. For a fixed number of total units, there is an optimal edit rate. For example, for *nU* = 20, edit rates of 0.3 ~ 0.7 are required to achieve a maximum encoding depth of two. By contrast, for *nU* = 500, edit rates of 0.02 ~ 0.03 could support new edits in each of 39 contiguous cell cycles. This analysis reveals that a extremely large number of coding units would be required for dense reconstruction of protracted cell lineages.

We also need to ensure sufficient number of encodable states (coding capacity) to confer co-derived cells with distinct codes. That is, especially crucial when mapping an exponentially increasing population of cells. Besides the number and possible assortment of cumulative edits, the number of levels (alleles) per unit contributes significantly to the coding capacity at a given depth. Fig. 1E exemplifies how various numbers of levels (*nL*) may impact the average capacity. First, this capacity is depth dependent since the number of edited units is depth dependent. Second, for *nL* = 1 the capacity converges to 1, since all fully edited codes are identical. Therefore, for systems with *nL* = 1, such as MEMOIR [3], care must be taken to stop editing while there is still enough encoding capacity. Third, when the number of units is reasonable, a small number of levels (*nL* > 1) per unit can encode a fairly large system. Even with *nL* = 2, the coding capacity exceeds 10^12^ for *nU* = 40 and reaches 10^150^ for *nU* = 500. Therefore, except for covering an enormous system with limited units, the number of levels is less critical than the number of units or the edit rate (as long as it is greater than 1).

Most biological systems are relatively small (e.g. 3.72 × 10^13^ cells per human body [4]) compared to the coding capacity of hundreds of units. However, a large caveat is that cellular production could involve rather protracted linear cell lineages (e.g. a typical neural stem cell in the tiny *Drosophila* brain undergoes ~ 100 serial cell cycles). Given the above modeling on editing dynamics, we need to further explore how to maximize the depth of coverage with a minimal number of coding units.

### Robust and efficient encoding of protracted cell lineages

From modeling a simple scheme of cumulative editing (Fig. 1C), we can determine the average number of new edits needed to obtain (with 99.9% probability) at least one new edit per cell cycle. Ensuring at least one new edit per cell cycle will enables us to differentiate each step of lineage depth. For a small rate (*r* < 0.1), we calculate the average number of expected new edits needed to be seven (*nlog*(1 − *r*) ~ *nr* ≤ *log*(0.001) ~ −7). Therefore, using seven as the minimal number of new edits per cell cycle, we can determine how many total coding units are needed (for a given editing rate) to record an entire lineage with 100 consecutive cell cycles. The optimal editing rate for recording 100 serial cell cycles is 0.01 with total required units of 1893 (Fig.2A middle). We define the efficiency of this scenario as 37%, derived as the ratio of the total units minimally required to cover the entire lineage (700) over the actual required number of units (1893). Lowering the editing rate reduces the efficiency because the majority of units are left unedited (Fig.2A left). By contrast, higher editing rates rapidly consume excessive units and therefore would require even more astonishing numbers of total units to sustain new edits throughout 100 serial cell cycles (Fig.2A right).

**Figure 2:**
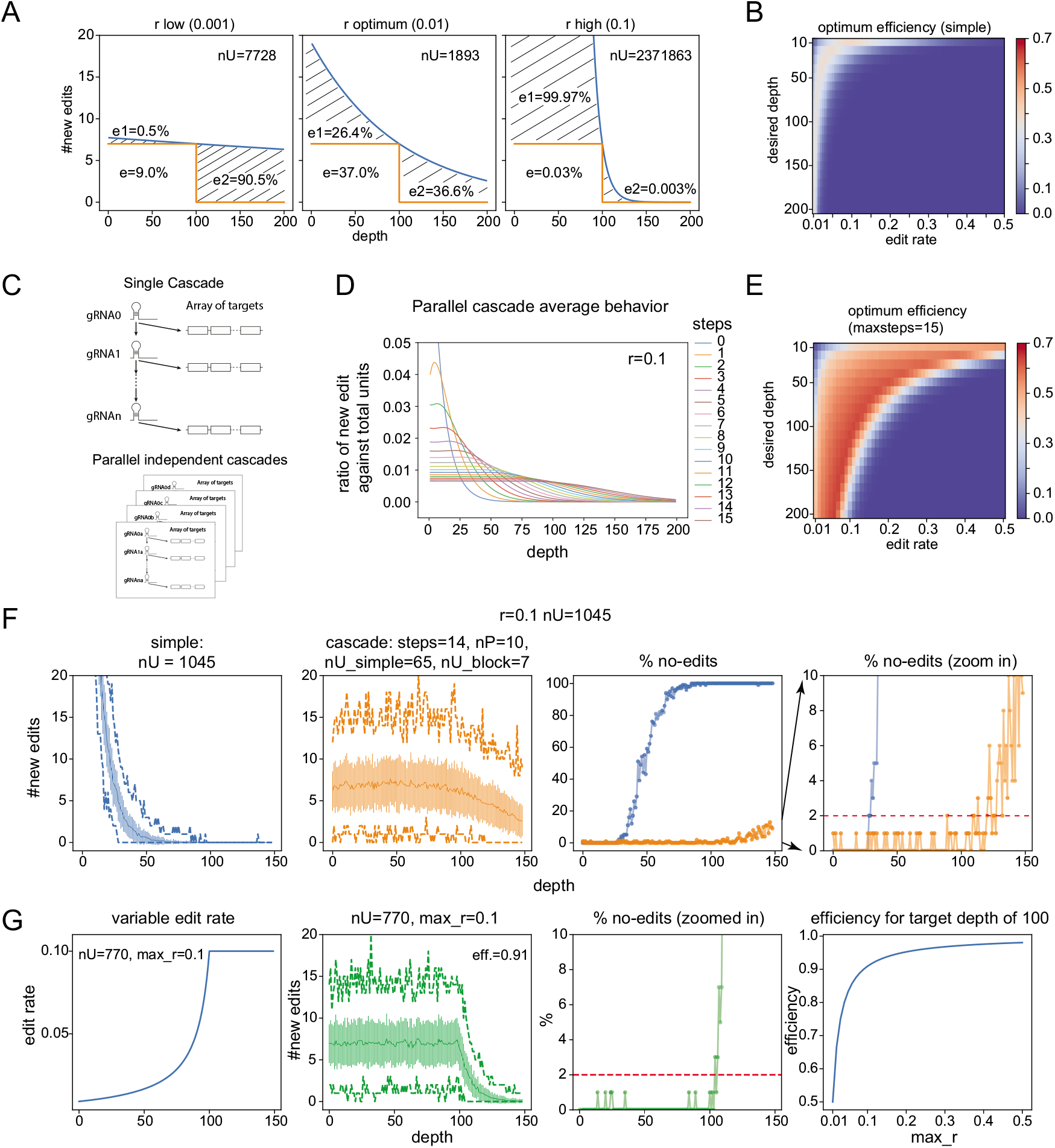
Dynamics of encoding. (A) Schematics showing the efficiency e for three cases of edit rates for simple exponential encoding process. The shadowed areas indicate wasted units. In the case of low edit rate (left panel), the waste resides mainly in the depth larger than the desired depth (100). In the case of high edit rate (right panel), the waste resides mainly in the initial phase. (B) Heatmap showing the efficiencies for various edit rates and desired depths for simple exponential encoding process. (C) Schematics showing a single cascade encoding process (top) and parallel cascades (bottom). (D) Average new edits (divided by total number of units) for parallel cascades with various number of cascade steps. Average is calculated as sum of infinite parallel cascades. (E) Similar to panel B, efficiencies of the cascade process for various edit rates and desired depths. (F) (Left two panels) Average number of new edits (middle solid lines), standard deviations (vertical bars) and minimum and maximum (dashed lines) of the number of new edits at each depth are shown for simple exponential case (1st panel) and for cascade process (2nd panel). Cascade process is realized with 10 parallel cascades of 14 steps with 7 target units per block and a single block of 65 pre-activated targets, constituting total of 1045 targets. (Righ two panels) Percentages of occurrences of no-edit at each depth are shown for simple (blue) and cascade (orange) processes. (G) (1st panel) An example of optimum variable edit rates when *max_r* = 0.1. (2nd panel) Similar to left two panels of F. Average, standard deviation, minimum and maximum of the new edits corresponding to the variable edit rate in the 1st panel are shown. (3rd panel) Zoomed in view of the percentage of no-edit. (4th panel) Optimum efficiency for different *max_r* in the case of desired depth of 100.

We calculated the coding efficiency across a range of lineage depth and a wide range of editing rates (Fig.2B). We learn that the maximal efficiency for recording serial cell cycles with simple cumulative editing can barely reach 40%. In general, for a desired depth of *d*, the optimum edit rate happens at *r* = 1/*d* with efficiency of (1 − 1/*d*)^*d*−1^, which is < 0.4 for *d* > 6. Moreover, the coding efficiency is very sensitive to the editing rate (Fig.2B), which is often uncontrollable. Modifying the editing dynamics would establish a more robust barcoding systems with higher efficiency and therefore enable dense reconstruction of protracted cell lineages.

To achieve the best possible coding efficiency with stochastic edits, we must maintain the average number of expected new edits around seven throughout the entire lineage. We envision two strategies to render the average number of expected new edits relatively stable across serial cell cycles, one controlling availability of units and the other controlling the edit rate. The first strategy involves subsets of the barcodes to be edited in successive bouts, where gRNAs targeting particular coding units are activated in a step-wise manner. The second strategy depends on varying the editing rate across serial cell cycles to compensate for the otherwise exponential reduction in editable units along the lineage depth. A promoter of an endogenous gene with proper expression dynamics maybe used for gRNA promoter.

Sequentially supplying gRNAs to edit separate pools of CRISPR targets would naturally increase the depth of coverage, as it would eliminate the problem of over editing at the earlier phase. The challenge is how to automate this process and distribute the overall coding capacity as evenly as possibly across an entire lineage. Recently, we developed a pair of novel strategies that utilize Cas9 as genetic switches.

CaSSA [5] permits gRNA-dependent gene reconstitution. Based on the CaSSA system, CLADES (Garcia-Marques et al., in preparation) provides a strategy for gRNA-dependent gRNA reconstitution. Using this methodology, we can reconstitute distinct gRNAs in a preprogrammed sequence as a cascade. Like other CRISPR edits, the Cas9-dependent progression of the gRNA cascade is stochastic. This means that a single cascade would produce uneven editing throughout lineage development. However, by combining multiple independent (parallel) cascades, the overall progression of the serial editing can be smoothened (Fig.2C, D).

To aid modeling cascade-driven serial edits, we assume comparable rates for gRNA reconstitution and editing of discrete gRNA-specific targets. In addition, we assume infinite parallel cascades (where mathematics is simpler) in the assessment of the best possible coding efficiency the cascade system can achieve. Plotted in Fig. 2D are the average edits per cell generation (shown as percentage of total units) for increasing numbers of cascade steps. Intriguingly, as the number of steps increase from 2 to 15, the distribution of edits across the depth is gradually flattened. This increases the efficiency of the unit usage. This flattened portion can readily extend beyond 100 serial cell cycles, enabling tracking of protracted lineages. Utilizing such plots, we can derive the highest achievable efficiency, with parallel gRNA cascades, for various combinations of editing rate and desired depth (Fig.2E). Compared to simple cumulative CRISPR edits, the cascade-based system not only increases the maximum efficiency from 0.37 to 0.70 but also greatly broaden the high-efficiency domain to a wide range of editing rates (Fig.2B vs. Fig.2E). As to practical applications using limited numbers of parallel cascades, we demonstrate via computer simulation that a system with 10 parallel 14-step cascades to drive editing of 1045 units in total can effectively cover 100 serial cell cycles with ~ 7 new edits per cell cycle. Without cascades, the depth of coverage for 1045 coding units drops from 100 to ~ 30 cell cycles. Taken together, our modeling demonstrates that driving CRISPR edits with parallel gRNA cascades can improve the coding efficiency by delivering a rather constant number of new edits per cell cycle up to a desired depth.

Another potential way to maintain a constant number of new edits per cell cycle is by increasing the editing rate along the depth. This increased editing rate can ensure new edits even as the number of unedited units drops. Strikingly, we can achieve a coding efficiency of 0.91 by setting the editing rate inversely proportional to the depth: *r_k_* = *ne*_0_/(*nU* − *k*\**ne*_0_), where *ne*_0_ is the desired number of edits per generation, which is 7 here. In such a system, the average number of new edits per cycle is kept at *ne*_0_. We further restrict *r_k_* from exceeding a defined maximum editing rate (*max_r*), which in Fig.2G is 0.1. The resulting curve of increasing editing rate mimics the ascending expression levels of some temporal genes (e.g. Syp) in the cycling neural stem cells. Therefore, we can possibly ramp up the editing rate by simply placing Cas9 expression under the control of a temporal gene promoter such as *Syp* [6]. In summary, modeling the dynamics of cumulative CRISPR edits has informed us how to encode protracted cell lineages with robust and efficient barcodes.

### Faithful reconstruction of densely encoded cell lineages

Now that we have modeled the best ways to perform CRISPR/Cas9 encoding, the next question is how best to decode a CRISPR/Cas9 coded lineage. Table 1 summarizes lineage reconstruction methods used in previous publications. Due to obvious similarity between lineage branching and evoluationary phylogeny, phylogenetic algorithms such as maximum parsimony[13] and neighbor joining[14] have been used with CRISPR/Cas9 coded datasets. Other groups have used a custom graph-based algorithm (LINNAEUS[9]), or hierarchical clustering together with a distance metric and linkage method ([12, 3]). Despite the variety of algorithms available to decode CRISPR/Cas9 encoded lineage data, no systematic comparison has yet been performed.

**Table 1:** Summary of previously employed CRISPR-based lineage reconstruction methods

| Assay Name | Tree Building Method | Comments | Ref |
| --- | --- | --- | --- |
| (sc)GESTALT | Maximum Parsimony by Camin- Sokal algorithm (using PHYLIP- mix, encoding each pattern as 0 or 1) | Since a pattern can span multiple targets, if each pattern is treated as a unit, then they may not be independent to each other. | [7, 8] |
| MEMOIR | Hierarchical clustering with average linkage (UPGMA), with custom tie breaking. Also NJ(Neighbor Joining) and Maximum Parsimony are used for simulation. | MEMOIR "scratch pad" only has two states (intact and mutated). This only works for limited depth range, since if all the scratch pad are mutated, they cannot be distinguished. | [3] |
| LINNAEUS | Custom algorithm (top-down, ordered by the abundance of scar among cells) | Uniqueness of scar, i.e. a same scar (pattern) does not happen as two independent events), is a critical assumption of the algorithm, which is often not met. | [9] |
| Molecular Recorder | Likelihood maximizing tree search with FNW (Frequency Normalized Weighting) of mutations to bias against more likely trees. | Belongs to the same category as Max. Parsimony in a sense that they are both forward tree search strategies maximizing some objective function (parsimony or likelihood). This class of algorithm is very slow. | [10] |
| Salvador-Martinez | Neighbor Joining using PAUP* and FastTree (for large samples) | Neighbor Joining is used over Maximal Parsimony due to speed issues. | [11] |
| Byungjin Hwang | Hierarchical clustering with Jaccard distance metric. | Uses L1 repeats as targets and base editor for encoding. Largest number of units (>200). ( First tried LINNAEUS method but switched to hierarchical method. ) | [12] |

We therefore set out to evaluate all of these algorithms. In addition, we also explored an extensive combinations of linkage methods and metrics for hierarchical clustering. For this purpose, we simulated a diverse set of ground truth trees as inputs (642 different trees), then codes of the leaf nodes are supplied to various algorithms for reconstruction, then errors of the outputs were calculated against the ground truth trees.

As we showed in the previous section, the number of new edits decreases rapidly as the depth increases, therefore we expect the reconstruction reliability to also be depth dependent. This means that the reconstruction error rate should be much smaller for an earlier depth. To evaluate reconstruction performances of the whole tree, several inter-tree distance metrics such as the Robinson-Foulds metric[15] or pair-wise shortest path difference[16] etc. have been previously used. Here, we adapted the Robinson-Foulds metric in a depth dependent manner for a much finer understanding of the reconstruction error (Fig.3A, see legend and methods for details).

**Figure 3:**
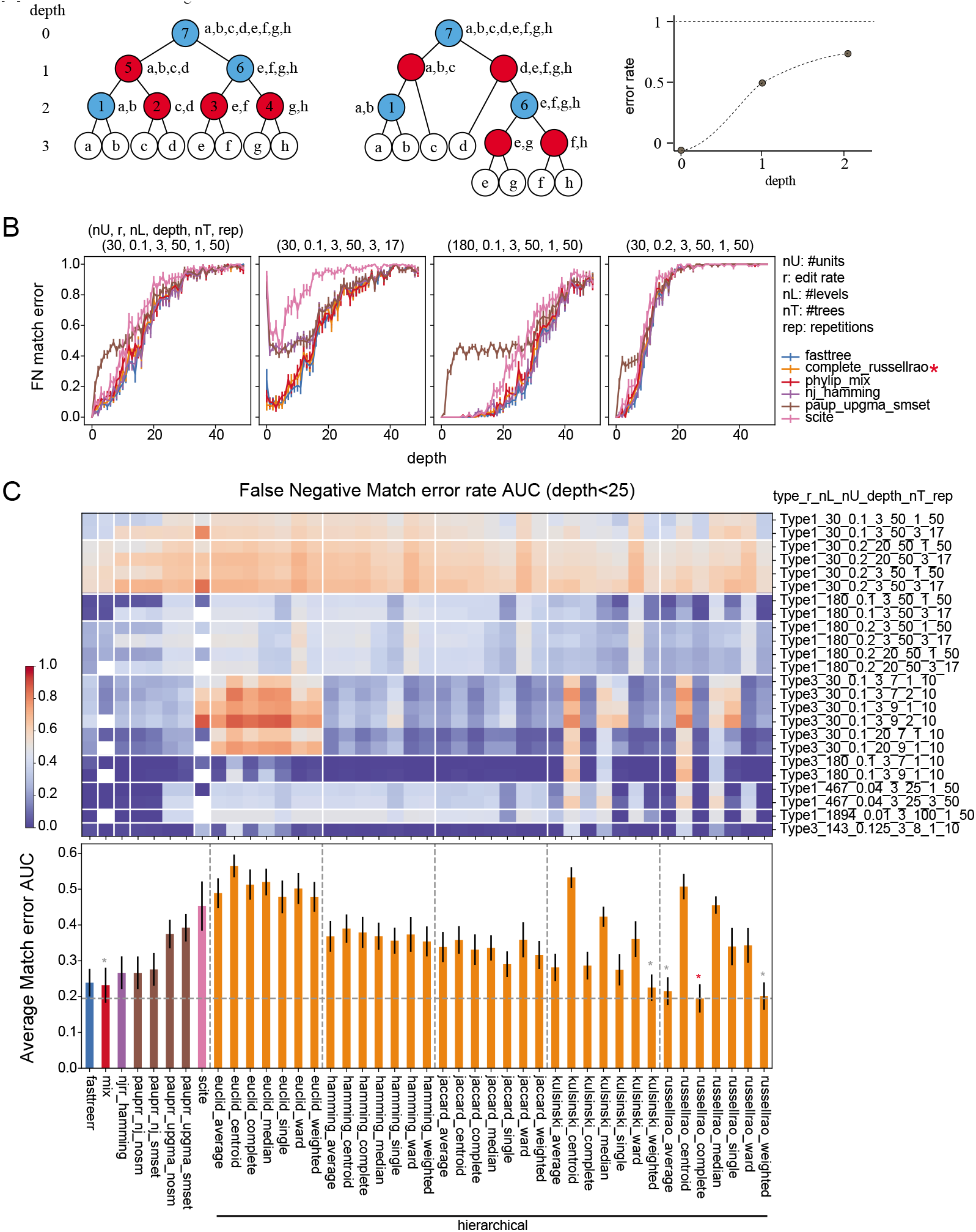
Performance evaluation of various reconstruction methods. (A) A simple example showing calculation of depth dependent error rate. The blue nodes in the original tree have matching nodes in the reconstructed tree but red nodes do not. At each depth in the original tree, the ratio of the number of the red nodes to the total number of nodes in that depth is the depth dependent error rate. (B) Depth dependent error rates for select conditions of inputs and reconstruction algorithms. The inputs shown here are Type1 (see method). Number of units (*nU*), edit rate (*r*), number of levels (*nL*) are as indicated. Number of trees (*nT*) indicate how many trees are combined and number of replicates (*rep*) indicate how many repeats are performed to generate the averaged traces. The errorbars indicate standard errors. Some conditions (such as linnaeus) does not show up which indicates that the execution took too long and terminated. (C) Heatmap of area under the curve (AUC) for depth <25 for all the conditions (inputs/algorithms) tested. The parameters for inputs are encoded as in panel B and shown in y-axis. The x-axis is common to next panel D and indicates the algorithms tested. For details on the inputs and algorithms see methods and supplements. (D) Averages of the AUC for all the input conditions. Error bar indicate standard error. For hierarchical clustering methods, linkage method (such as average, centroid) and distance metric (such as Euclid, hamming) combinations are indicated as (linkage)_(metric). Red star indicates the reconstruction method with the lowest error rate. Smaller gray stars indicate other methods statistically indistinguishable to the lowest case.

We simulated the CRISPR encoding process for a range of parameters (*nU,r,nL*) with different division modes (type 1: asymmetric division and type 3: symmetric division, see method for details). Some examples of optimal cases for simple exponential encoding were also include (bottom 4 cases in Fig.3C). In some cases, 3 trees (*nT* = 3) are combined to assess whether algorithms can successfully separate them. From the simulated trees, leaf nodes were extracted as inputs. Each input was then supplied to all of the surveyed algorithms and the reconstructions were evaluated against the ground truth. For the same set of parameters, this is repeated 50 times (17 when number of trees is 3) to account for the randomness of editing. Then the average depth dependent errors were calculated. Plotted in Fig.3B are a subset of conditions showing the average depth dependent error rates. As expected, earlier depths have smaller error rates. There are also clear performance differences between different algorithms. In some cases, (PHYLIP mix or SCITE), the execution took more than 20 min and so reconstruction was terminated and flagged as a failure (white boxes in Fig.3C). This was more than 2000 times the typical execution time of other algorithms (see Fig.Supp.7).

Fig.3C, D summarizes the performances of different algorithms with various inputs. For simplicity, the area under the curve (AUC) of the depth dependent error rate (with *depth* < 25) is shown. Fig.3C,D displays the false negative error rate (proportion of original nodes not reconstructed). We also assessed false positive error rate by calculating how many of the reconstructed nodes are absent from the original tree (Fig.Supp.4). In addition, we also calculated the false negative/positive error rate using more permissive error metrics (see Fig.Supp.5,6 and method). From these surveys, we found that hierarchical clustering with complete linkage and the Russell-Rao metric and FastTree2 which is based on Neighbor-Joining performs best over all.

For the metrics for binary numbers, such as hamming distance, we extended their definitions non-binary cases to distinguish levels (when *nL* > 1, see supplement). The Russell-Rao metric is one of such metric and with this extension, it is 1 minus the proportion of the number of edited units which have the same level between two codes. Therefore, we call this variation of RussellRao metric, “SharedEdits” (i.e. edits shared by two cells). Presumably these shared edits are inherited from a common ancestor.

### Why tracking SharedEdits outperforms more elaborate distance metrics?

In the previous section, we observed varied reconstruction performances between different hierarchical clustering methods. Hierarchical clustering iteratively merges a pair with the shortest distance and generates an ancestor node. Therefore, the choice of metric affects which pair is merged. To understand why the SharedEdits metric outperforms other metrics, we first factored out a common component, code duplication. Code duplication causes reconstruction errors regardless of the metric used. Code duplication between parent-offspring pairs occur either when all the units are edited (unit depletion) or when the remaining number of un-edited units is low causing no edits (Fig. 1C green line). Code duplication can also happen between non-parent-offspring pairs by chance but with a much lower frequency. Since no metric can distinguish cells with a same code, code duplication results in a tie and thus error (in half of the cases if tie breaking is random).

To remove the effect of code duplication, we modified the simulated trees to eliminate nodes with duplicated codes (see methods) and assessed the depth dependent error rate after reconstruction (Fig.4A orange dashed lines). This procedure dramatically reduced the error rates for the SharedEdits, Kulsinski metrics but not for Jaccard, Hamming or Euclid metrics. We investigated the places where errors occured and noticed that for the latter groups of metrics, there was a lot of precocious merging of shallower nodes. To remove the precocious merging, we then added a constraint that the depths of merging pairs be less than a given distance from the deepest remaining node during reconstruction (Fig.4A solid lines, see method). This constraint drastically reduced the error rate for the latter three metrics. This strongly indicate that the SharedEdits and Kulsinski metrics contain depth information suitable for proper reconstruction, whereas the latter three metrics do not.

**Figure 4:**
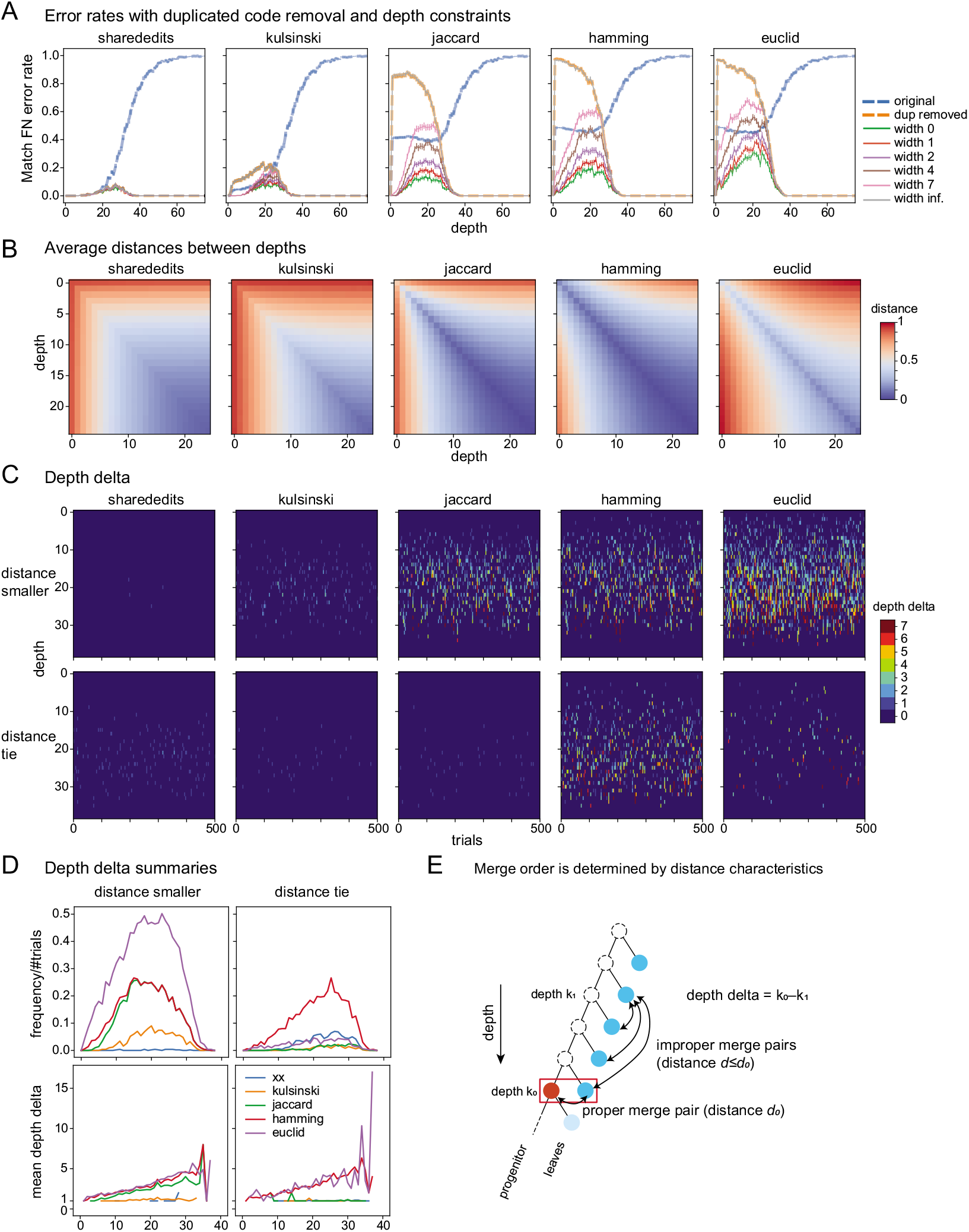
Why sharededits metric outperforms others. (A) Average depth dependent error rates for hierarchical clustering method with complete linkage for different distance metrics. Input is type 1, *nU* = 180, *nL* = 3, *r* = 0.1, *nT* = *1* and 500 replicates are used to calculate the average. Blue dashed lines show the error rates for original inputs. Orange dashed lines show the error rates after original inputs are modified to have no code duplication. The rest of the lines show the error rates with depth constraint (with depth width indicated) together with code duplication removal. (B) Pairwise average distances between nodes in different depths. Diagonals (*k*_0_, *k*_0_) are averages of distances between a progenitor and a leaf, (see panel E red and blue nodes at depth *k*_0_). Off-diagonals (*k*_0_, *k*_1_) are averages of minimum distances between a leaf (blue node at depth *k*_0_) and nodes in depth *k_0_* (both red and blue nodes at depth *k*_0_). (C) Heatmap showing the color-coded “depth delta” (see panel E) for each depth (y-axis) and replicate (x-axis). (D) (top row) Average frequencies of distance swap or tie across trials. (bottom row) Average depth delta when distance is swapped or tied. (E) Definition of depth delta.

To understand this in more detail, we plotted heatmaps showing the distances between nodes of different depths (Fig.4B). One noticeable difference between initial two and the latter three metrics is the diagonal part, distance to self. In fact, the SharedEdit and Kulsinski metrics are not full distance metrics but pseudo distance metrics in that the self-distances are not necessarily zero. This turned out to be a prerequisite for a more relevant feature of these metrics, that the distance between nodes is essentially a the depth of the shallower node and monotonically decreases with depth. This is indicated in the inverted L-shape patterns of the distance matrices, though this is approximately so for Kulsinski metric, as can be seen from the distortion of the inverted L-shape patterns in Fig.4B. Because of this property, the SharedEdits and Kulsinski metrics would calculate that a larger distance for a pair with shallower nodes than the proper merge pair, thus preventing precocious merging of shallower nodes. With the latter three metrics, despite the minimal distances of the proper merge pairs (diagonal elements in Fig.4B), the distortion surrounding the diagonal elements are likely to cause inappropriate precocious pairs with distances that are equal to or even smaller than the proper pair.

To see whether this is in fact the case, we calculate the occurrences of precocious pairs being either smaller (swap) or equal to (tie) the distance of the proper merge pair (see Fig.4C, E). We separated the cases of distance swap and tie because they contribute differently to the error rates. Swaps always lead to errors, however, in the case of ties, the proper merge pair can be selected by chance. The number of swaps with the SharedEdits metric was extremely low (Fig.4D top rows). While the Kulsinski metric had fewer ties than SharedEdits, it had much larger swaps resulting in potentially more errors. As expected, the other three metrics had much higher occurrences of swaps (and in some cases ties). Moreover, when we quantified the distance of the improper merge pair (depth delta, see Fig.4E), the latter three metrics had much larger values (Fig.4C, Fig.4D bottom rows), whereas SharedEdits and Kulsinski always had a depth delta less than 3 (Fig.4C, D). These indicate that overall distance characteristics of the SharedEdits and Kulsinski metrics (Fig.4B) prevented the frequent precocious merging of shallower nodes. In the cases where precocious merging occurred, it was restricted to a local range. Between the SharedEdits and Kulsinski metrics, we found that distance swaps occurred more frequently in Kulsinski and the depth deltas were larger (Fig.4D), thus SharedEdits performs better than Kulsinski. When swap/tie frequencies were calculated for depth-delta<2 (Fig.Supp.8), the frequencies closely resembled that of residual error in Fig.4A, indicating we’ve captured the source of the errors.

Since SharedEdits errors are almost always due to distance tie, we tried several tie-breaking methods. Unfortunately, none of them made noticeable improvement (Fig.Supp.9).

In summary, the SharedEdit metric outperforms other metrics because it encodes depth information in a manner that enables the reconstruction to proceed from the deepest pair to shallower pairs in a proper order.

## Discussion

CRISPR permits multiplex genome editing, which greatly facilitates engineering of intricate genetic barcodes for tracking cell lineages. Through theoretical modeling and computer simulation, we demonstrate the unparalleled promise of CRISPR-derived genetic barcodes in mapping protracted cell lineages with a single-cell-division resolution. We learned that the patterns of edits shared across barcodes best depict cell lineage relations. While in hindsight this observation seems rather obvious, it has been overlooked in previous reconstructions of CRISPR-coded cell lineages. Appraising cell relatedness based on shared barcode edits permits the best reconstruction of cell phylogeny. However, the lineage depth one can track remains severely limited for standard encoding process especially when the total unit number is small and high editing rates are needed to ensure robust barcode editing in early cell cycles. Our modeling demonstrated that it is possible to slow down the depletion of available coding units without sacrificing robust barcode editing. We can accomplish this by utilizing parallel gRNA cascades enabled by CLADES (Garcia-Marques et al., in preparation) or by using variable edit rates. With these encoding methods, we estimate that we need approximately one thousand independent coding units to densely track stem-cell-type lineages across 100 cell generations. Given CRISPR’s enormous versatility, it should be possible to develop further sophisticated genetic barcodes for comprehensive analysis of whole-organism development.

### Engineering CRISPR barcodes in vivo

Creating genetic barcodes for tracking cell lineages requires DNA sequences editings specifically in cycling cells. For continuous tracking, the early DNA edits should not be erased by later edits. For CRISPR encoding, we can provide units containing gRNA target sequences clustered together in a transgene(s) [17, 9, 7] or we can target widely distributed endogenous sequences, such as transposon-related repeats [12]. As to tandem targets, one major concern is interference from nearby targets. Two double strand breaks within the barcode could result in deletion of all intervening targets including those already edited [7]. Such complications would compromise overall coding capacity and could disrupt hierarchical clustering (due to the loss of various earlier edits). Targeting endogenous repeats, by contrast, would better ensure independent edits. Encouragingly, one recent study has demonstrated the feasibility of editing endogenous L1 elements in cultured cell lineage tracking [12]. However, utilizing endogenous sites requires detailed characterization of the genome to identify ideal targets (in terms of editing efficiency and edit retrievability) that also play no role in normal development. Given the above pros and cons, the possibility of dispersing exogenous gRNA targets (e.g. placing them sparsely on a bacterial artificial chromosome) should also be explored as a way to avoid inter-target deletions. Another solution would be to create diverse barcodes via DNA base editing (e.g. use of Cas9-deaminase) rather than through repair of double-strand DNA breaks. In the absence of nuclease activity, one should be able to maintain independent edits among tandemly packed gRNA targets. However, one may need to increase the number of coding units drastically to compensate for the lower editing rates observed with existing Cas9 base editors.

The great versatility in the control of Cas9 further enhances the power of CRISPR in creating genetic barcodes. First, one should be able to modulate the editing rates by altering Cas9 expression levels. Moreover, for efficient barcoding of protracted cell lineages, Cas9 activity could possibly be dynamically controlled to achieve robust, yet minimal editing in each cell cycle. Second, one can express Cas9 in specific spatiotemporal patterns for tracking specific cell lineages or specific developmental stages. Further, it is important to restrict Cas9 to cycling cells to prevent post-mitotic editing, which would be non-informative and may overshadow precursor-derived edits especially in long-lived extant cells (e.g. neurons). Finally, gRNAs govern the target specificity of Cas9 actions. The virtually infinite specificity of gRNAs endows an unlimited multiplex power, which can be exploited to build further sophisticated genetic barcodes (see next). In brief, we see it feasible to record any dynamic biological activities across cell generations using CRISPR-derived genetic barcodes.

### Robust and efficient encoding

Robust barcoding requires occurrence of new edits in every cell cycle. Fewer coding units are needed to ensure robust barcoding for the higher editing rate. However, the higher the editing rate, the faster the available coding units would be depleted, thus preventing deep encoding. This dilemma makes tracking protracted stem-cell-type lineages extremely challenging. Given a hundred coding units, we need an editing rate of ~0.1 to achieve robust barcoding in early depth. But at a rate of 0.1, a hundred total coding units could not sustain robust barcoding beyond 10 cell generations (Fig.1D). At an editing rate of 0.1, to continuously track 100 cell generations (as needed for mapping most *Drosophila* neuronal lineages), we would need a daunting number (>2 million, Fig.4A) of coding units. Notably, we can drastically reduce the needed number of coding units to around two thousand by simply decreasing the editing rate to ~0.01. However, most of the units are still not efficiently utilized (Fig.4A). To improve barcoding efficiency, we should instead modulate the availability of coding units or edit rates across the depth of cell lineages. For maximal efficiency, we should use up all coding units and distribute the edited units randomly and evenly throughout the entire length of each lineage.

Strikingly, using parallel gRNA cascades to drive the editing, we can consume most coding units and distribute the edits rather evenly throughout a protracted lineage. Reserving separate pools of coding units for editing at different developmental times has been practiced manually by injecting a second gRNA set at a later stage of zebrafish development [7]. Thanks to CLADES (Garcia-Marques et al., in preparation) which allows serial reconstitution of multiple gRNA variants, we can now automate the process of supplying distinct gRNAs in a cascade to serially edit different subsets of coding units. We found that infinite parallel cascades mathematically produces a flat consumption of codes across depth (Fig.4D), enabling an efficient encoding process compared to the simple exponential case (Fig.4E). We also showed that this idealized behavior of infinite parallelism is essentially reproducible with a rather small number (10) of parallel cascades (Fig.4F).

We also showed that controlling editing rate can lead to very robust and efficient encoding system (Fig.4G). Even though the control of edit rate is not perfect, the efficiency is expected to be very high. The editing rate may be controllable by driving Cas9 gRNA under endogenous gene promoters which mimic the desired dynamics (such as *Syp* [6]) or under small molecule inducible promoters and manipulating the concentration of the inducer molecules. The change in the cell cycle duration can also be utilized. For example, *Drosophila* neuroblasts divide quickly in the early phase but the division becomes slow in the later phase of development. If Cas9 gRNA edit rate is constant per duration, then the change of cell cycle duration effectively increases the edit rate per division[10]. The exact configuration which may be able to achieve desired edit rate change should be a subject of future studies.

Taking all factors into consideration, we propose a cascade barcoding system for tracking *Drosophila* brain cell lineages with ~1000 coding units in total. This elaborate system ensures occurrence of 7 or more new edits per cell cycle (98% of the time) in all cell cycles, and consumption of >70% of the 1000 units by the end of 100 serial cell generations. Such cascade barcoding systems, driven by targeted Cas9 induction, would allow us to efficiently reconstruct cell phylogeny with single-cell-cycle resolution for any complex tissue.

### A better cell lineage reconstruction method

Various reconstruction methods have been utilized for CRISPR-based cell lineage codes (Table 1). Here, we explored the performances of these methods as well as other combinations of metrics and linkage methods in hierarchical clustering (Fig.3C). Notably, we found hierarchical clustering with a previously unused metric outperforms all other methods (Fig.3D). This metric, which we call SharedEdits, is based on the number of common edits.

We found that the reason the SharedEdits metric outperforms other metrics is because it contains the depth information in a way appropriate for reconstructing the tree in a proper order (Fig.4). The SharedEdits distance of a pair is essentially determined by the number of edits the common ancestor of the pair has. This helps to prevent precocious merging of nodes in shallower levels and promotes the appropriate bottom-up reconstruction of cell lineages. Other metrics (Jaccard, Hamming and Euclid) do not possess this property, and by imposing restriction in allowed depth during reconstruction, a drastic proportion of the reconstruction errors (> 50%) can be corrected (Fig.4A 3rd-5th panels, orange vs. green lines). The Kulsinski metric has a similar property to SharedEdits, but depth constraint can still improve the error rate (Fig.4A 2nd panel). This indicates that the way it contains the depth information is still not ideal. In contrast, the SharedEdits metric error rates did not improve upon depth constraint (Fig.4A 1st panel), indicating that the manner SharedEdits contains the depth information is near ideal.

No reconstruction method can solve the ambiguities associated with code duplication. Thus, the errors caused by code duplication set the limit in the reconstruction error rates. By artificially removing code duplication events, we found that the reconstruction using SharedEdits produces very small error rates (Fig.4A 1st panel orange line). This indicates that the reconstruction method based on SharedEdits is very close to optimum.

Since the residual errors (errors without code duplication) for the SharedEdits metric are mainly due to distance tie (Fig.4C, D), we have tried to improve it further by incorporating other metrics or factors (such as number of leaves under each node) to break the tie. These efforts produced little to no improvements (data not shown). We have not yet tried to combine other methods such as Nearest Neighbor Interchange which incorporate scores from larger structures or later steps.

### Additional needs for further sophisticated algorithms

For practical applications, we have tested the robustness of our optimized tree-building method in multiple aspects, including variable editing rates and unequal choice of edit outcomes, as well as substantial cell loss. We found that it performs robustly as long as the barcoding is sufficient and intact. However, the program in its current state is vulnerable to random loss of various coding units, which can happen in the code retrieval process (e.g. single cell genomic PCR and sequencing). This weakness argues again for the importance of having integrated barcodes that can be readily retrieved as a whole, thus protecting against random loss of code. Nonetheless, there are clearly unmet needs, including recovery of missing code and the scalability in processing big data sets, which we hope to address in the future with additional sophisticated algorithms.

In summary, we have found a better method for reconstructing CRISPR-coded cell lineages than previously used. This method is based on the metric using the number of shared Cas9 edits. We have further demonstrated alternative encoding strategies in tracking every cell cycle across numerous cell generations with high efficiency. Such versatile genetic barcoding empowered by CRISPR technology will revolutionize how we study biological organisms.

## Materials and Methods

### Depth dependent error rate

There are several distance metrics that quantifies similarity between trees. Of these, Robinson-Foulds distance (RF distance)[15] is arguably the most frequently used. For two rooted trees, RF distance counts 0 for internal nodes which exist in both and 1 for which exist only in one tree (internal node match is determined by the set of leaves under the node). We extended this metric to depend on depth in the following way. We first designate one of the trees (usually the original simulated tree) as base tree and flag each internal nodes of the base tree depending on whether it has a matching internal node in the other tree (usually the reconstructed tree). Then for each depth of the base tree, we calculate the proportion of the nodes which do not have a matching node in the other tree (Fig.3A). This set of numbers is a natural extension of RF distance to depend on depth, albeit being normalized per depth. We call these numbers “match” error rates. If we use original tree as the base, then this error rate is false negative error rate since it counts the original nodes not present in the reconstructed tree. If we use reconstructed tree as the base, the it becomes false positive error rate since it counts the reconstructed nodes not in the original tree.

There are other ways to extend whole tree metrics to depend on depth. For example, we can restrict the target of matching to be in the same depth in the other tree in the above example, or we can first restrict trees to subtrees down to a depth and then calculate whole tree distance between these subtrees as depth dependent distance. We have tried these metrics as well with similar qualitative results (data not shown). We have also extended another tree metric, *pairwise cell shortest-path distance* [16] in to a depth dependent metric using subtrees described above. This also resulted in qualitatively similar results (data not shown).

The above match error rate is highly sensitive since it flags an internal node as an error node even if only one leaf contained in the node is wrongly assigned out of many other leaves. To devise an error metric which can distinguish 1 wrong assignment out of 100 versus 10 wrong assignments out of 100, we search the node in the non-base tree which has the maximal similarity to the base node in terms of leaf set similarity measured by Jaccard index. We then assign for each depth average max Jaccard similarities for nodes in the depth. We call this depth dependent error metric jaccard error.

### Lineage tree simulations

Here we tested two types of lineage trees. One increases linearly with depth (type1), and the other increases exponentially with depth (type3). A type 1 progenitor asymmetrically divides and generates one type1 and one ganglion mother cell (GMC) which turns into two terminal cells (leaf nodes). We simulated CRISPR encoding to happen once during a cell cycle before cell division, which resulted in the two offsprings of a GMC sharing one code, so we can simply say type1 progenitor generates another type1 progenitor node, one GMC and one leaf node. A type 3 progenitor symmetrically divides and generates two type 3 progenitors. At the desired depth, we stop the division and simply set those nodes without offspring to be leaf nodes. For the inputs to the reconstruction algorithms, only codes from leaf nodes are supplied.

### Reconstruction algorithms

We tested reconstruction algorithms previously used in CRISPR-based lineage tracing publications. Of these Neighbor-Joining (NJ) [14] and maximum parisimony [13] are from traditional phylogeny field. We used the same software packages and settings as much as possible. In some cases, we also tested other implementation or settings as well. For NJ, we used PAUP* [18] with several different setting (see table below) and FastTree [19] as these are used in the previous CRISPR-based lineage tracing publications. We also tested *nj* function from skbio.tree Python package (http://scikit-bio.org) as an alternative implementation since it accepts input as distance matrix. For maximum parsimony, we used mix from PHYLIP package [20] which was used in GESTALT publications [7, 8]. We also tested a software package (scite [21]) from tumor lineage field as an example of algorithms which utilize a stochastic search algorithm such as MCMC method. For method used in LINNAEUS paper [9], we modified the author’s R script to accept input in a more convenient format but used as is otherwise. For hierarchical clustering, we tested combinations of 7 linkage methods (average, centroid, complete, median, single, ward and weighted) and 5 distance metrics (euclid, hamming, jaccard, kulsinski, russellrao). Distance metrics are chosen to include popular ones (euclid, hamming) as well as ones used in previous publications (jaccard) and top performing metrics (russellrao and kulsinski) from preliminary survey of more extensive set of metrics. For binary metrics (hamming, jaccard, kulsinski and russellrao), we modified them to distinguish edit outcome levels (see supplement for exact definition). The *linkage* function in scipy.cluster.hierarchy Python package (http://www.scipy.org) is used to perform the hierarchical clustering. Most of the standalone programs output in nexus or newick formats. We used DendroPy (http://dendropy.org) to parse these outputs. See supplemental material for exact details on algorithm used in Fig.3.

### Removal of code duplication

To filter out the effect of code duplication, we first compared all the pairs of nodes using hamming distance and detected duplicated pairs as entries with 0. We then removed nodes with duplicated code with larger depth. When the node being removed is not a leaf node, child nodes of the node are reconnected to the parent node of the node being removed. In the case of trees used in Fig.4 (type 1, *nU* = 180, *nL* = 3,*r* = 0.1, *nT* = 1), this removal procedure shrunk the tree depth to < 40. The exact topology of the trees depend on the instance (trial).

### Depth constraint in reconstruction

Depth constraint during reconstruction is enforced so that the minimum estimated depth of the next merging pair should be within the current maximum estimated depth minus 2. For leaf nodes, depths are assigned using the original tree information. For internal nodes with matching node in original tree, again depth information from the original tree is used. For internal nodes without matching node, depth is inferred as minimum depth of the nodes under the node minus 1.

## Supplementary Materials

### Cascade efficiency with various parameter ranges

**Figure Supp.1:**
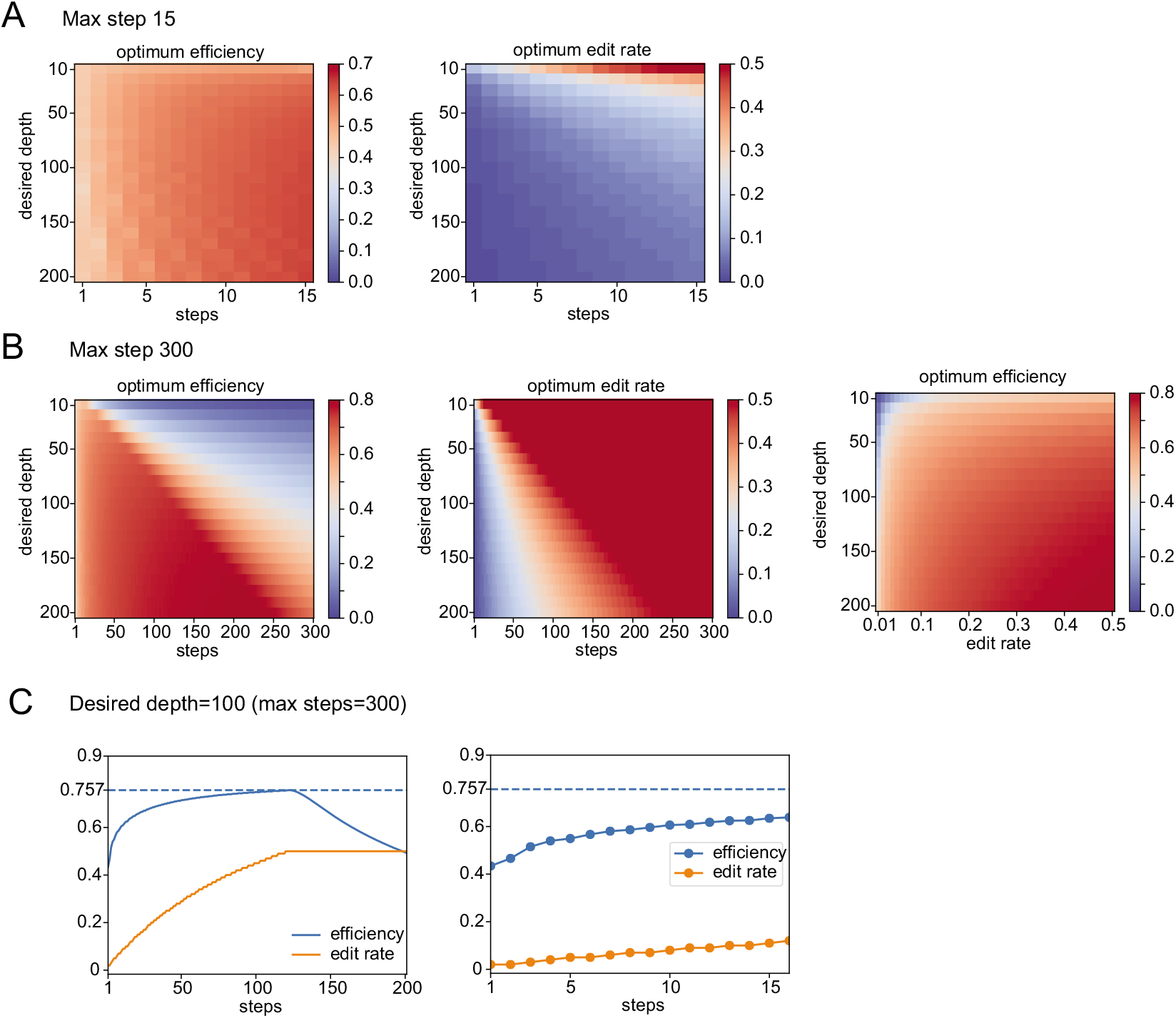
Parameter ranges for CLADE cascade system. To calculate optimum caso of cascade system, we first need to specify the range of cascade steps (15 in A and 300 in B) and the range of edit rates (0.01 to 0.5). Within these fixed ranges of cascade steps and edit rates, we can find the optimal combination of these for desired depth to achieve maximal efficiency. Panels shown in A and B shows relationships between these parameters. Panel C shows the optimal efficiency and edit rate for fixed steps in the case of desired depth of 100.

### Variable edit rate robustness against linear approximation

**Figure Supp.2:**
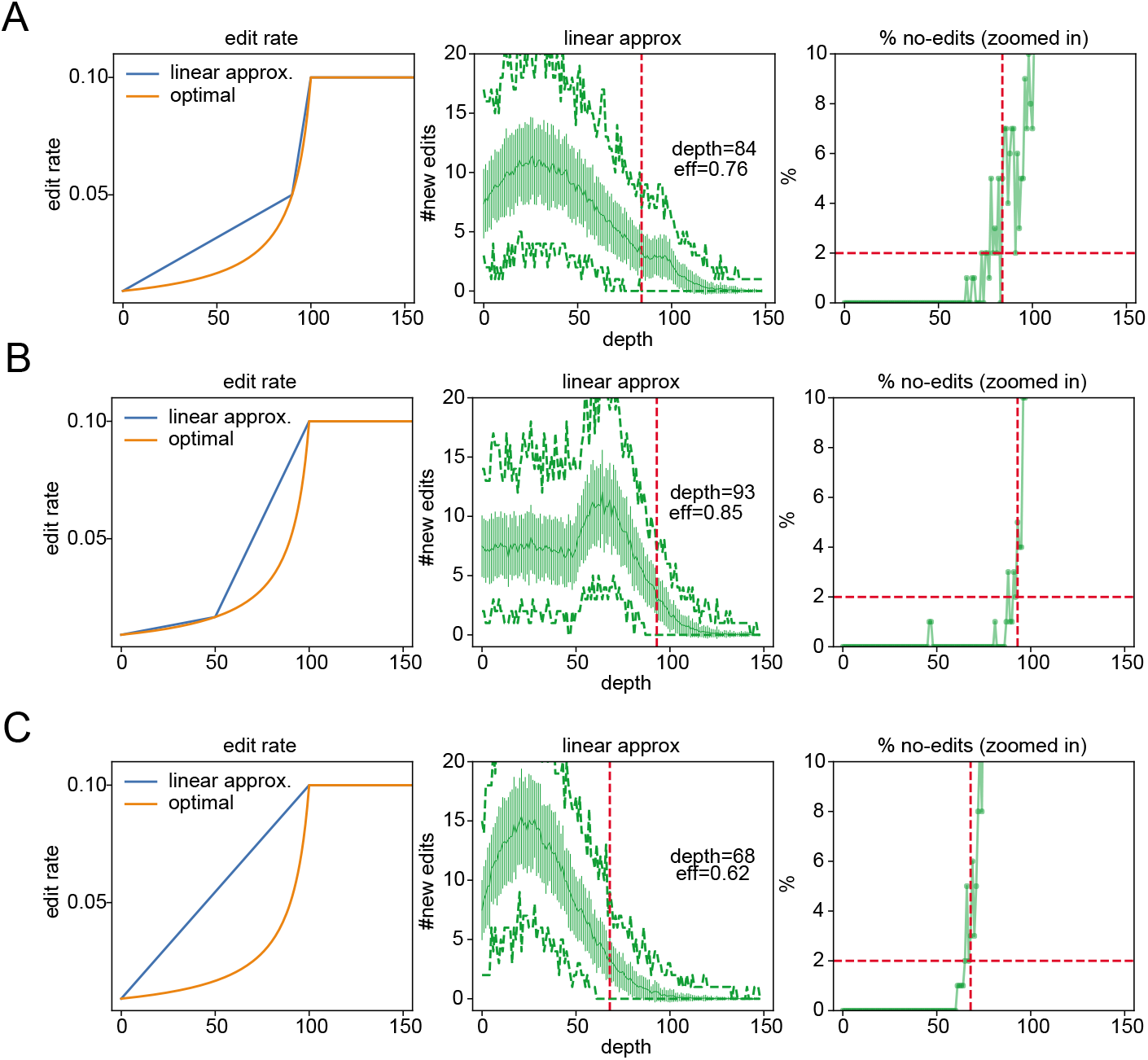
Piece-wise linear approximation of variable edit rate system. Various pioce-wiso linear approximation of variable edit rate system are shown. When optimal variable edit rate is approximated by piece-wise linear rates, the reachable depth and efficiency are reduced. The efficiencies are. however, still much higher than those achievable by simple exponential encoding.

### Variable edit rate robustness against noise

**Figure Supp.3:**
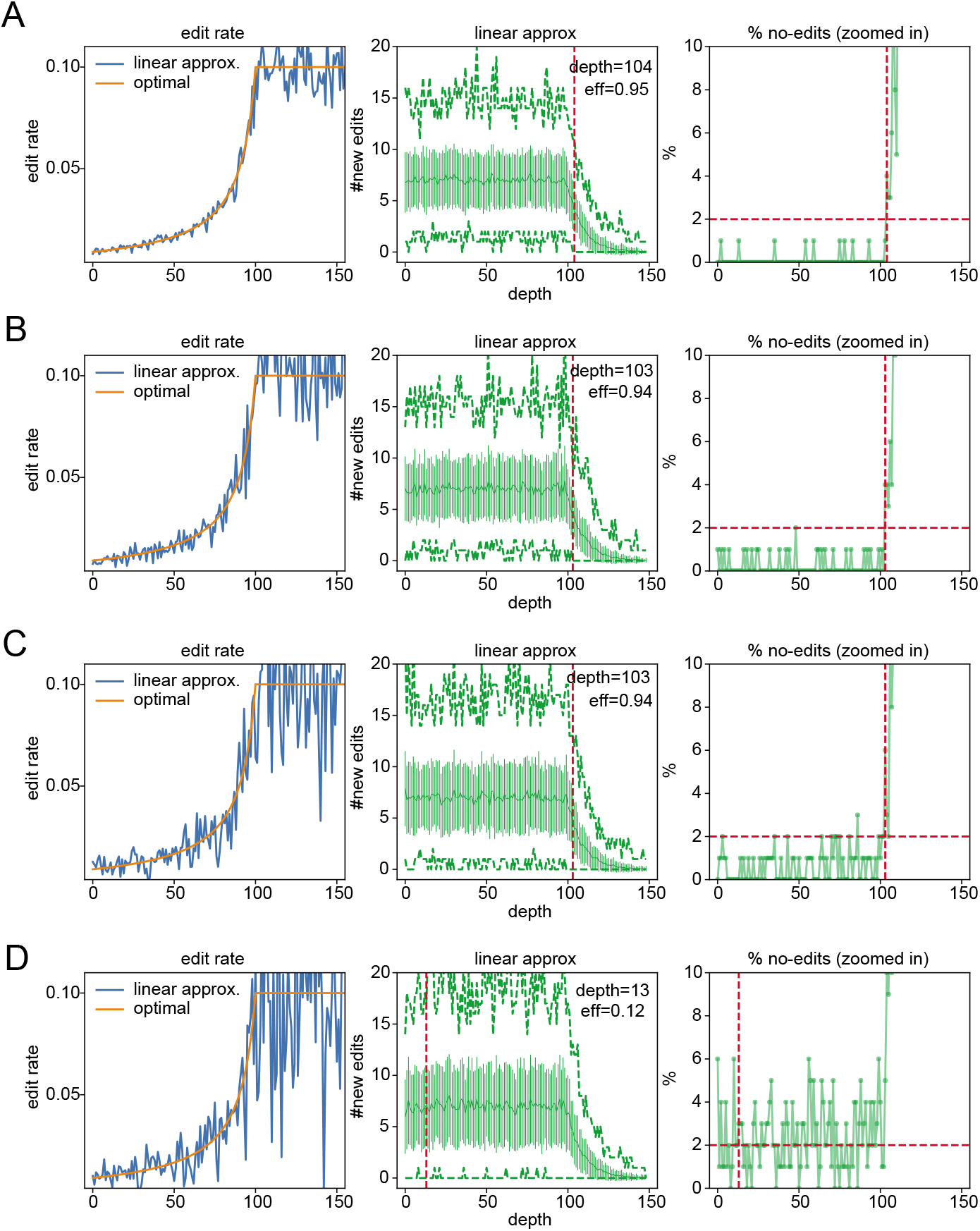
Effect of noise in variable edit rate system. Various noise level proportional to the rate are introduced to the variable edit rate system (A: 10%. B:20%. C:30% and D:40%). The system is robust against up to 30% noise.

### Performance evaluation: Match error (False positive rates)

**Figure Supp.4:**
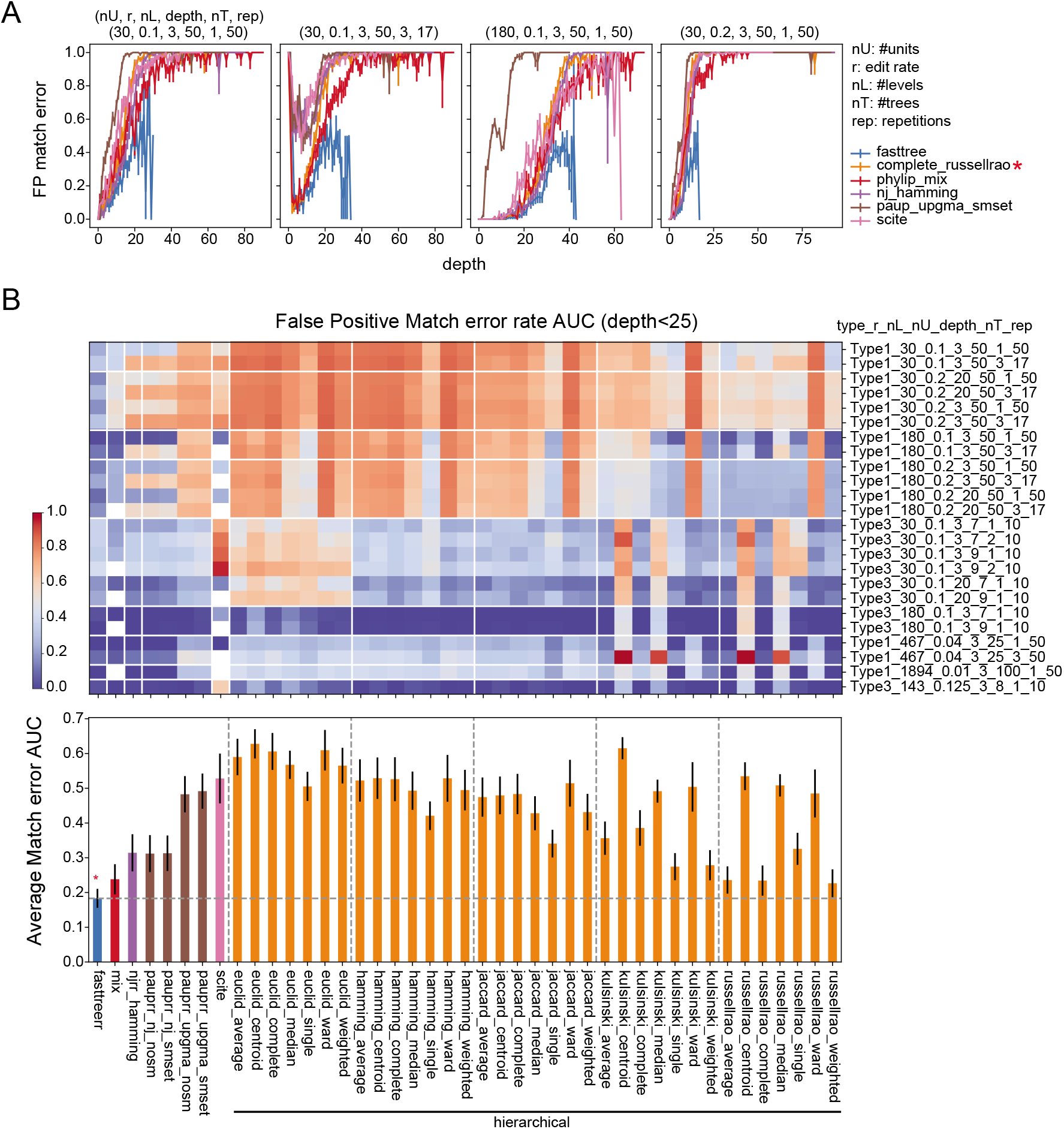
False positive error rates measured by match error. Similar to Fig.3 but false positive error rates (proportion of reconstructed nodes not present in the original tree) are shown.

### Performance evaluation: Jaccard error (False negative rate)

**Figure Supp.5:**
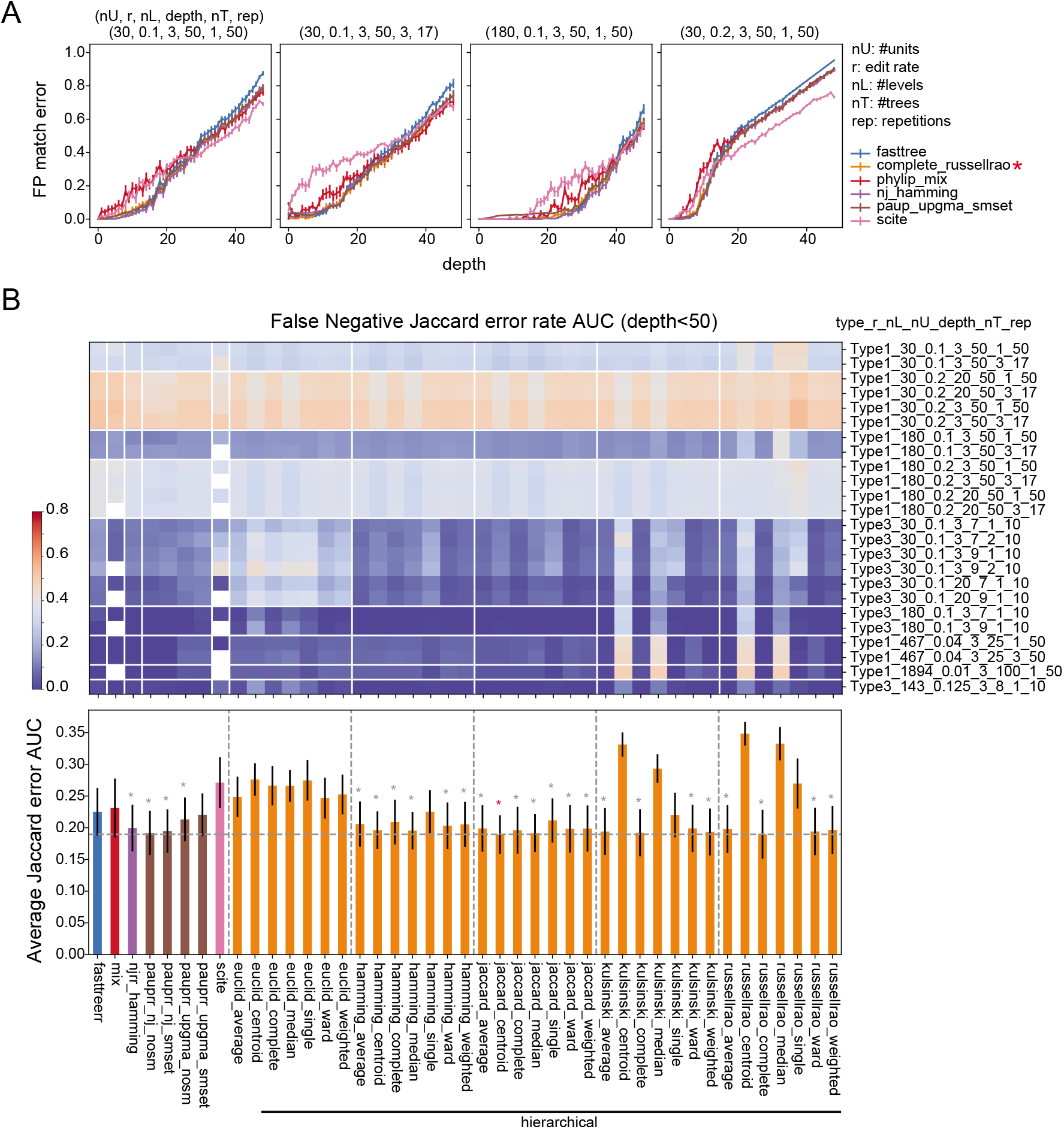
False negative error rates measured by Jaccard error. Similar to Fig.3 but false negative error rates using Jaccard error (see method) are shown.

### Performance evaluation: Jaccard error (False positive rate)

**Figure Supp.6:**
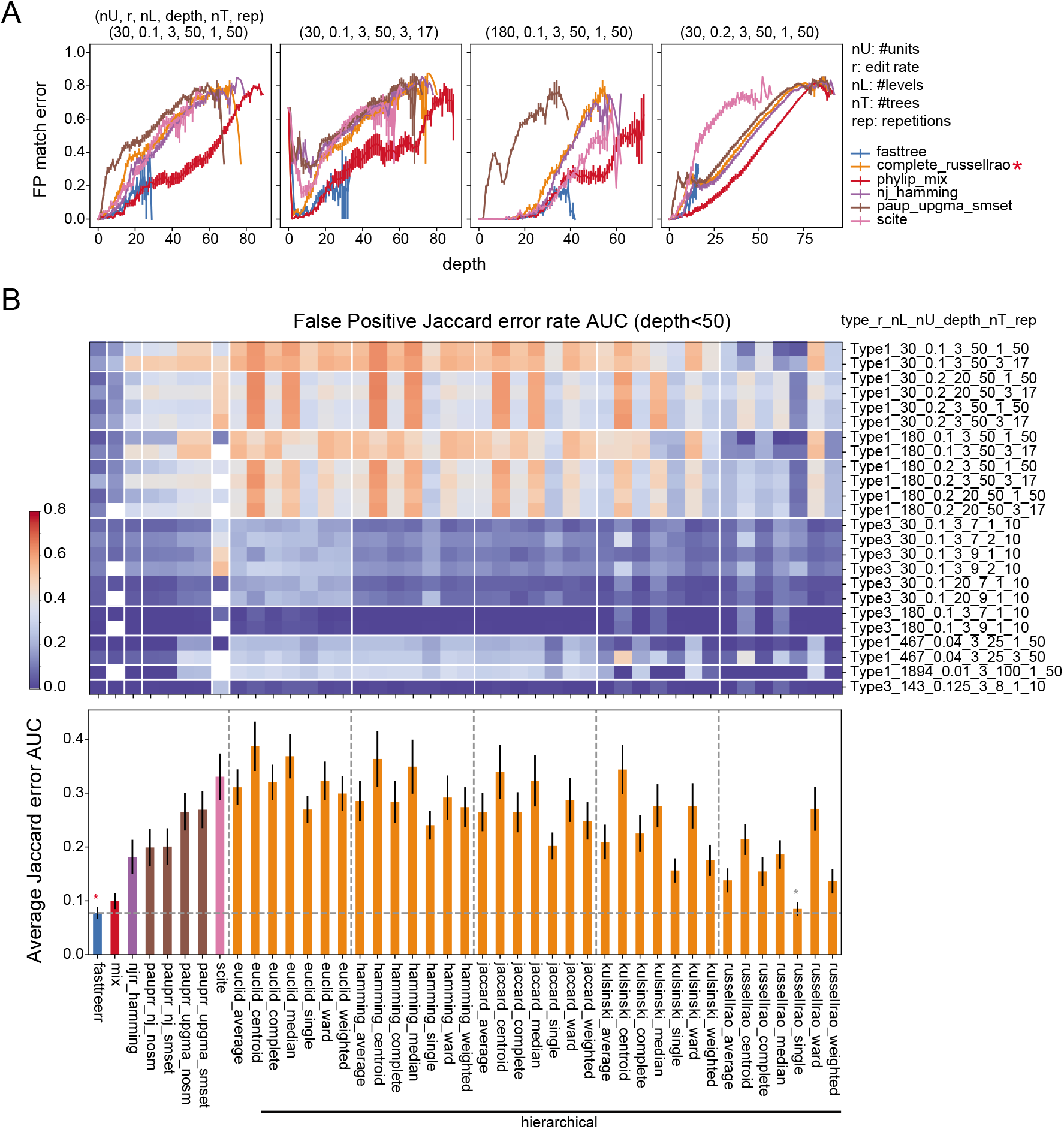
False positive error rates measured by Jaccard error. Similar to Fig.3 but false pasitive error rates using Jaccard error (see method) are shown.

### Performance evaluation: execution time

**Figure Supp.7:**
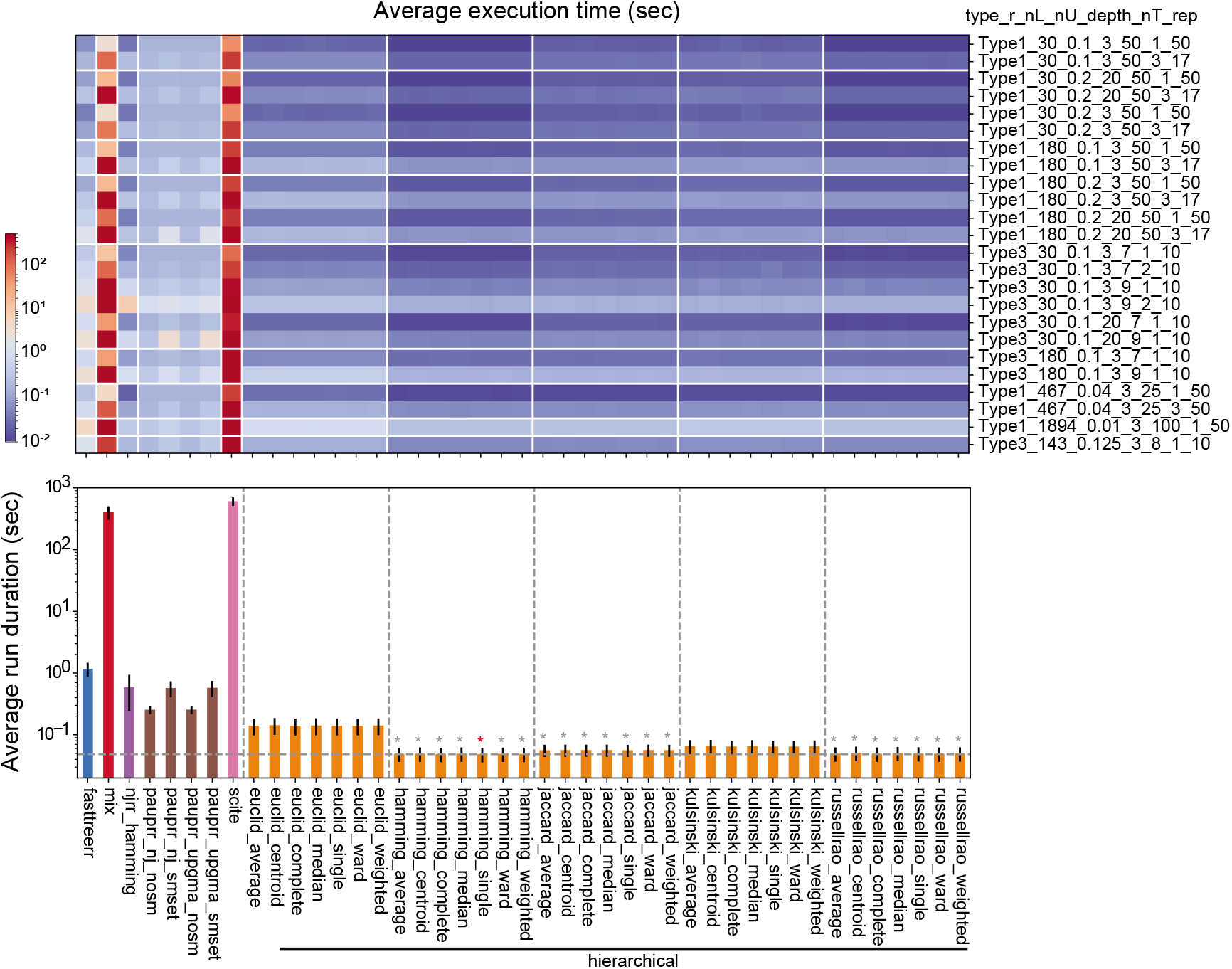
Program execution duration summary. Summary of execution duration for each program/input combinations. Note the scale for duration (sec) is in logarithm.

### Type1 error source supplements

**Figure Supp.8:**
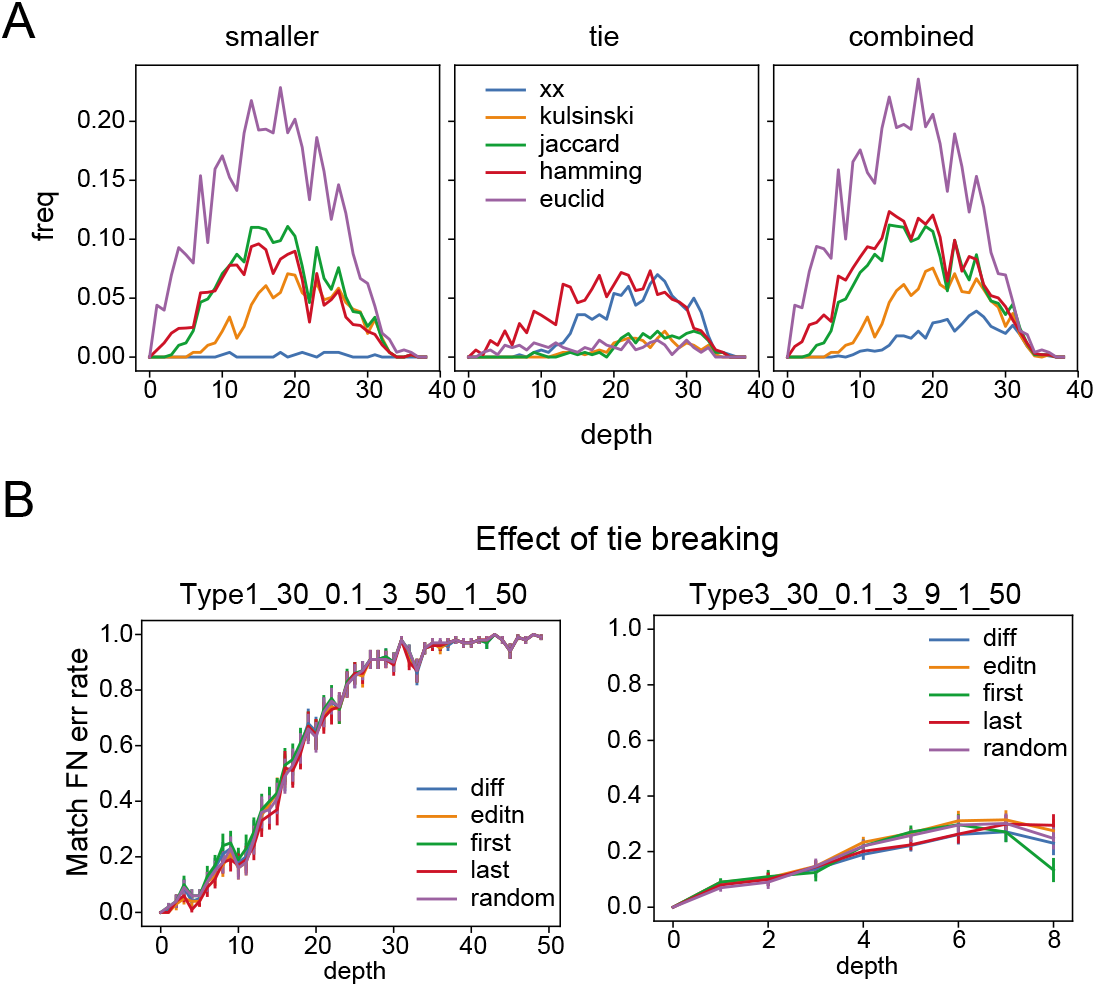
Type1 error source supplement. (A) Distance swap/tie frequencies for depth delta<2. The third panel shows combined frequency calculated as *swap* + *tie*/2. (B) Effect of various tie-breaking, “diff” uses hamming distance to break the tie. “editn” calculates how many units are edited and chooses one with the largest number of edits, “first”.”last” chooses the first (or last) of the ties, “random” randomly chooses one from ties.

### Error source for type3 tree

**Figure Supp.9:**
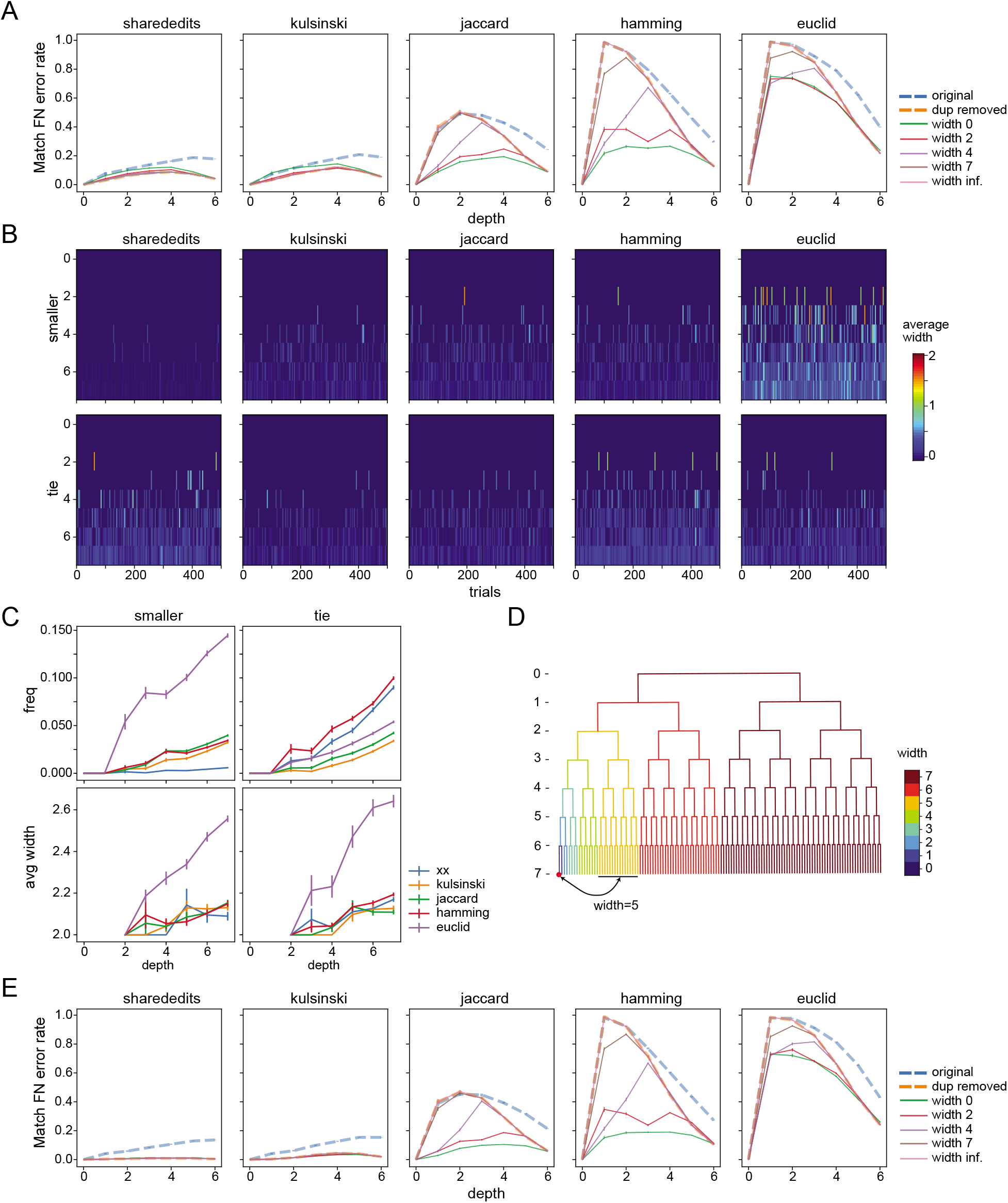
Performance comparison of metrics: type3 case. Similar to Fig.4 but calculated against type3 case (*nU* = 30, *L* = 3, *r* = 0.1, *depth* = 7, *nt* = 1, *nrep* = 500). Instead of depth-delta, we calculated “width” (defined in panel D) for each depth. Panel E shows the case of L=20 instead of L=3. Shared edits and kulsinski metrics improve with larger number of levels but three other metrics do not.

### Details of algorithm execution

**Table Supp.1:** Settings for algorithms in Fig.3

| Abbreviation in Fig.3 | Package | Metric | Comments |
| --- | --- | --- | --- |
| fasttree | FastTree2 | n/a | Levels are encoded as nucleotide string (ATCG; if levels are more than 3, then multiple characters are used) and codes are then converted to FASTA format and supplied to standalone fasttree executable, with option “-nt -gtr”. Unedited code (root) is include in the input and then used to reorder nodes in the output (re-rooting). |
| (linkage)_(distance) | scipy | euclid,<br>hamming,<br>jaccard,<br>russellrao,<br>kulsinski | Input to the scipy hierarchical clustering method is a distance matrix and which linkage method (average, centroid, complete, median, single, ward, weighted) to use. |
| linnaeus | LINNAEUS | n/a | Unit and level combination is treated as one entity (called scar) and its presence is indicated for each cell as an input. Linnaeus produces an output tree with scars as nodes. We converted this to a tree with cells as nodes by replacing scars with matching cells. |
| phylip_mix | PHYLIP | n/a | Unit and level combination is treated as one entity and its presence/absence is represented as 1 or 0. Thus, codes of length $n_u$ are converted to strings of 0 and 1 of length $n_u n_l$. Weights are calculated as log ratio of counts of each unit/level combination to the maximum counts and encoded in a character array. Then mix program is executed with ifile,P,W,2,3,Y,wfile options, where ifile and wfile are the names of input and weight files. |
| nj_hamming | scikit-bio | hamming | skbio.tree.nj function takes distance matrix as an input, hamming distance is used. Re-rooted as in the case of FastTree. |
| nj_russellrao | scikit-bio | russellrao | Similar to above, russellrao distance is used. |
| paup_nj_nosm | PAUP* | n/a | Codes are converted as FASTA as in fasttree as inputs. NJ method is specified. No user stepmatrix is specified. Since NJ is unrooted, re-rooting is done as in FastTree. |
| paup_nj_smset | PAUP* | n/a | As in above, but stepmatrix is specified with log of inverse frequency as the weights. (Same as in [11]). |
| paup_upgma_nosm | PAUP* | n/a | As in paup_nj_nosm but upgma method is used instead of nj. |
| paup_upgma_smset | PAUP* | n/a | As in paup_nj_smset but upgma method is used instead of nj. |
| scite | SCITE | n/a | Codes are converted to strings of 0,1 as in phylip_mix and supplied as an input to scite and executed with options “-i ifile -n nm -m nc -r 2 -l 400001 -fd 1e-2 -ad 1e-1 -transpose”, where ifile is input file name, nm is number of mutations, nc is number of cells |

### Probabilistic Model of CRISPR Editing

The number of edited units, *e*_1_, from total of *n_u_* units with per cell division edit rate of *r* follows binomial distribution which we denote *B*(*n_u_, r*). That is, probability of getting *e*_1_ edits is 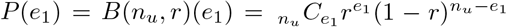. The complement of this, i.e. the number of un-edited units, *n*_1_ = *n_u_* − *e*_1_, follows *B*(*n_u_,s*), where *s* = 1 − *r*. In the second round of edits, getting *e*_2_ new edits again follows binomial distribution: *B*(*n*_1_, *r*), but with a different parameter *n*_1_ which depends on the outcome of the 1st round. The combined distribution is a multinomial distribution:

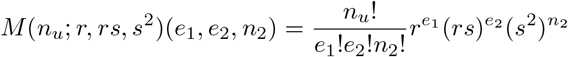

Where *n*_2_ = *n_u_* − *e*_1_ − *e*_2_ is the number of unedited units after 2nd division. In general, after *k* cell division, probability of getting *e*_1_, *e*_2_,…,*e_k_* edits at each division is a multinomial distribution:

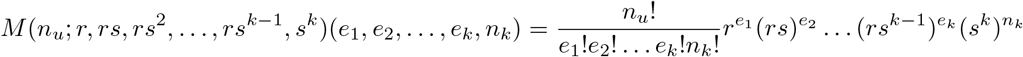

Where 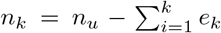 is the number of unedited units after *k*-th division (Fig.1B). Because of the property of the multinomial distribution (next section, Lemma 3), the total number of edited units after *k* cell division, 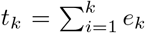, follows binomial distribution *B*(*n_u_,r* + *rs* + *rs*^2^ +…+ *rs*^*k*−1^) = *B*(*n_u_*, 1 − *s^k^*). Similarly, the number of unedited units *n_k_* follows binomial distribution *B*(*n_u_, s^k^*) with standard deviation 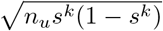, (Fig.1C). We will define effective editing rate at depth *k* as *r_k_* = 1 − *s^k^* and its compliment as *s_k_* = 1 − *r_k_* = *s^k^*.

### Properties of Multinomial Distribution

#### Lemma 1.

*Sum of two random variables x*_12_ = *x*_1_ + *x*_2_ *in a multinomial distribution M*(*n;r*_1_, *r*_2_,…,*r_k_*) *follows multinomial distribution M*(*n; r*_1_ + *r*_2_, *r*_3_,…,*r_k_*).

*Proof*. Using binomial expansion:

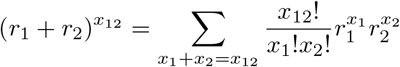

it follows probability for *x*_12_ which is a sum of *x*_1_, *x*_2_ over *x*_1_ + *x*_2_ = *x*_12_ is:

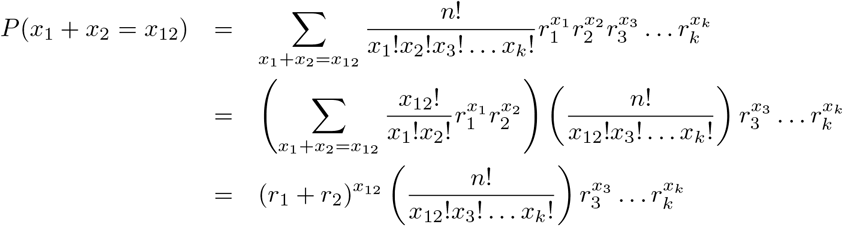

thus, *x*_12_ (and *x*_3_,…,*x_k_*) follows *M*(*n; r*_1_ + *r*_2_, *r*_3_,…,*r_k_*).

#### Lemma 2.

*Sum of random variables in multinomial distribution again follows multinomial distribution whose rate is the sum of rates of summed variables*.

*Proof*. Apply lemma 1 multiple times.

#### Lemma 3.

*Multinomial distribution integrated over all but one variable is a binomial distribution*:

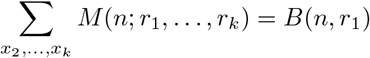

*Proof*. Applying lemma 2 on variables *x*_2_,…,*x_k_* will yield the desired result since a multinomial distribution of degree 2 is a binomial distribution.

#### Lemma 4.

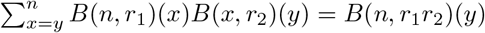

*Proof*. Using,

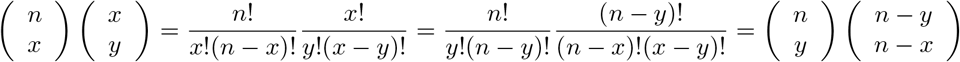

and replacing *n* − *x* = *x*′, where *x*′: 0 → *n* − *y*, since *x*: *y* → *n*,

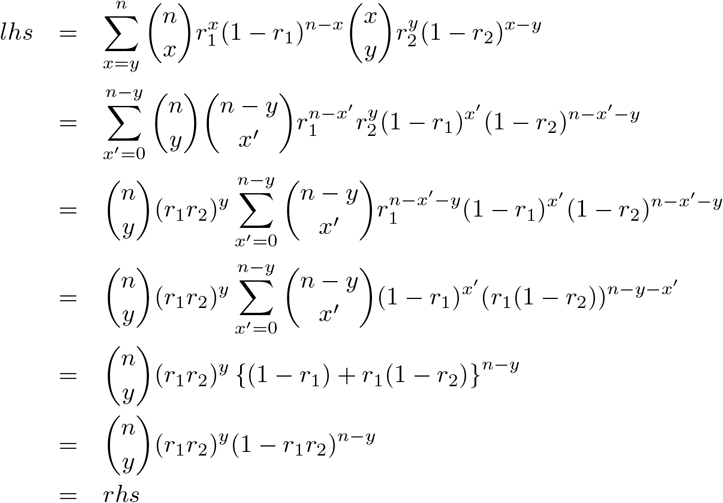

### Distance Metrics

**Figure Supp.10:**
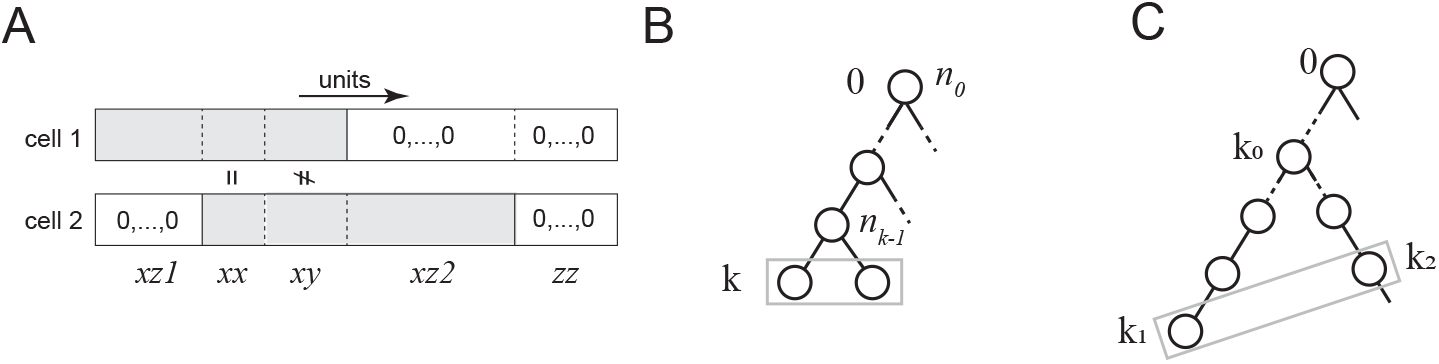
5 random variables for code match between two cells The number of units which share the same edits between two cells is denoted *xx*. The number of units which are edited in both cells but not same is denoted *xy*. The numbers of units edited only in one cell are denoted *xz*1 and *xz*2. The number of units unedited in both cells is denoted *zz*.

We denoted the number of outcomes of editing in a unit as *n_l_*(excluding unedited state) and assumes the probability of choosing any outcome is the same, i.e.: *p* = 1/*nL*. When considering two lineage codes, there are 5 basic random variables which quantifies the difference between them (Fig.Supp1A). These are: *X_zz_*, which is the number of unit unedited in both, *X*_xz1_ mid *X*_*xz*2_, which are the number of units edited in one but not the other, *X_xx_*, which is the number of units edited in both and having the same edit outcome, and *X_xy_*, which is the number of units edited in both but with different outcomes. For single round of edits, these variables sum up to the total number of units, *nU*(= *n*_0_), and follow multinomial distribution with rates *rs* (for *X*_*xz*1_ and *X*_*xz*2_), *r*^2^*p* (for *X_xx_*), *r*^2^*q* (for *X_xy_*), and *s*^2^(for *X_zz_*), where *q* =1 − *p*. When comparing two sibling codes, which are at the depth of *k* (Fig.Supp1B), we need to take into account the distribution of the number of unedited units at depth *k* − 1: *n*_*k*−1_. For example, probability of getting *X_xy_*, is:

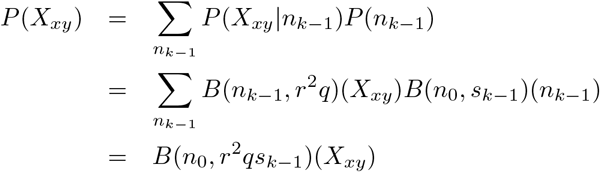

using Lemma 4 in the previous section. That is, rate is simply multiplied by *s*^*k*−1^. Similarly for *X*_*xz*1_, *X*_*xz*2_ and *X_zz_*. For *X_xx_*, we need to take into account shared code up to the previous depth *k* − 1,which is *t*_*k*−1_ ~ *B*(*n*_0_, *r*_*k*−1_) = *B*(*n*_0_, 1 − *s*^*k*−1^), so the rate becomes 1 − *s*^*k*−1^ + *r*^2^*ps*^*k*−1^. In general, if two codes are separated as in Fig.Supp1C, then, the rates for *X_xx_,X_xy_, X*_*xz*1_, *X*_*xz*2_, *X_zz_* are

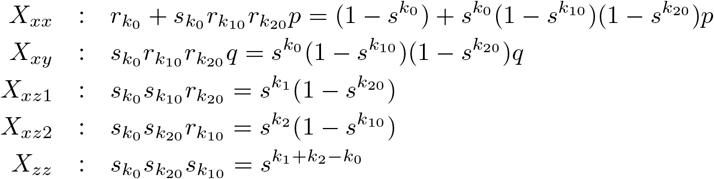

where *k*_10_ = *k*_1_ − *k*_0_ and *k*_20_ = *k*_2_ − *k*_0_.

Using these variables, we can also define the (extended) distance metrics between two lineage codes as follows:

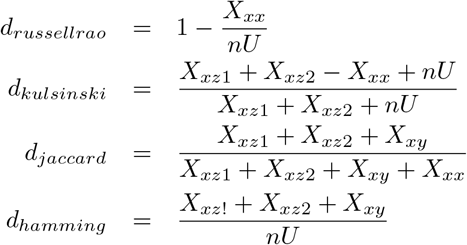

